# Reward-predictive cues elicit maladaptive reward seeking in adolescent rats

**DOI:** 10.1101/2020.06.17.157644

**Authors:** Andrew T. Marshall, Nigel T. Maidment, Sean B. Ostlund

## Abstract

Impulsive behavior during adolescence may stem from a developmental imbalance between motivational and impulse control systems, producing greater urges to pursue reward and weakened capacities to inhibit such actions. Here, we developed a Pavlovian-instrumental transfer (PIT) protocol to assay rats’ ability to suppress cue-motivated reward seeking based on changes in reward expectancy. Traditionally, PIT studies focus on how reward-predictive cues motivate instrumental reward-seeking behavior (lever pressing). However, cues signaling imminent reward delivery also elicit countervailing focal-search responses (food-cup approach). We first examined how reward expectancy (cue-reward probability) influences expression of these competing behaviors. Adult male rats increased rates of lever pressing when presented with cues signaling lower probabilities of reward but focused their activity at the food cup on trials with cues that signaled higher probabilities of reward. We then compared adolescent and adult male rats in their responsivity to cues signaling different reward probabilities. In contrast to adults, adolescent rats did not flexibly adjust their pattern of responding based on the expected likelihood of reward delivery but increased their rate of lever pressing for both weak and strong cues. These findings indicate that impulse control over cue-motivated behavior is fundamentally dysregulated during adolescence, providing a model for studying neurobiological mechanisms of adolescent impulsivity.

## 1. Introduction

Adolescents have a penchant for impulsive and risky behavior. This willingness to act in the face of uncertainty may be adaptive, prompting them to assert newfound control over their actions and establish independence from caregivers (Casey et al., 2008; Spear, 2000). But adolescents are also prone to pathological forms of reward-seeking behavior, such as binge drinking, unprotected sex, and reckless driving (Steinberg et al., 2008). Advancing our understanding of the mechanisms of adolescent impulsive behavior through well-controlled animal research may ultimately inform the development of new approaches to combat these and related public health concerns.

It is widely believed that the rise in impulsive behavior during adolescence is driven in part by developmental changes in emotional and motivational systems which result in intense urges to pursue reward (Casey et al., 2008; Spear, 2000; Steinberg, 2010). Consistent with this view, previous studies have shown that adolescent rats exhibit heightened palatable food intake and hedonic reactions to palatable food stimuli, as well as an increased willingness to exert effort to consume food reward (Friemel et al., 2010; Marshall et al., 2017a; Schneider et al., 2015; Stolyarova and Izquierdo, 2015; Wilmouth and Spear, 2009).

Reward-predictive cues can be powerful triggers of impulsive reward-seeking actions, which raises the possibility that adolescents may be particularly vulnerable to their motivational influence. Interestingly, animal studies that have taken up this issue provide relatively little support for this view. Initial studies using the Pavlovian conditioned approach paradigm found that adolescent rats are less – not more – likely than adults to approach and interact with reward-predictive cues (Anderson and Spear, 2011; Doremus-Fitzwater and Spear, 2011), an index of incentive motivation known as sign-tracking. However, more recent studies have found that adolescent rats do not differ from adults (Anderson et al., 2013) and may even show elevated levels of sign-tracking behavior (DeAngeli et al., 2017) under some conditions (e.g., social isolation and food deprivation), pointing to the need for further research on this question.

The Pavlovian-to-instrumental transfer (PIT) paradigm has also been used to probe developmental changes in cue-motivated behavior. The PIT task is unique in that it focuses on the tendency for reward-paired cues to elicit independently trained, instrumental reward-seeking responses such as lever pressing (Dickinson et al., 2000; Estes, 1948, 1943; Rescorla and Solomon, 1967). Because the cue and the lever-press response are never paired during training, the cue’s ability to trigger lever pressing provides a relatively pure and unambiguous measure of its incentive motivational influence. Using this approach, Naneix et al. (2012) found that cue-elicited reward seeking did not significantly differ between adolescent and adult rats, once again suggesting that the motivational influence of reward-paired cues is not simply more intense during adolescence.

However, developmental delays in impulse control are also thought to play an essential role in adolescent impulsive behavior, weakening the ability to suppress maladaptive reward-seeking actions that may be triggered by a potentially overactive motivational system (Casey et al., 2008; Ernst et al., 2006; Steinberg, 2004). This view is supported by reports that adolescent rats engage in heightened levels of reward seeking in situations where such behavior is unnecessary or counterproductive (Andrzejewski et al., 2011; Burton and Fletcher, 2012). Given such findings, we hypothesized that adolescent rats may be more susceptible to the motivational influence of reward-paired cues under conditions in which this impulse to seek out reward is normally suppressed.

The current study sought to disentangle the contributions of *incentive motivation* and *impulse control* to cue-motivated reward seeking and determine how these processes are altered during adolescence, relative to adulthood. Our strategy involved developing a novel PIT task to probe cue-motivated reward seeking under varying levels of response conflict (and therefore different impulse control requirements). In a recent report (Marshall and Ostlund, 2018), we found that the tendency for reward-paired cues to motivate lever-press performance was replaced by food-cup approach behavior when cue conditions signaled imminent reward delivery. This suggests that adult rats are normally able to suppress their motivational impulse to engage in exploratory reward seeking (e.g., lever pressing) when reward is strongly expected so that they can instead engage in more situationally advantageous focal search behavior (e.g., food-cup approach). However, it remains unclear how adolescent rats would respond under such conditions. We reasoned that if there is a developmental imbalance between motivational and impulse control systems during adolescence, then adolescent rats should have particular difficulty inhibiting the maladaptive impulse to lever press in the presence of strong reward-predictive cues.

## 2. Methods

### 2.1. Experiment 1

This experiment investigated the influence of expected reward probability on cue-elicited lever pressing and food-cup approach behavior in adult rats, allowing us to establish appropriate conditions for characterizing adolescent behavior in Experiment 2.

#### 2.1.1. Animals

Thirty experimentally naïve male Long Evans rats (Envigo) were used in this experiment. They arrived at the facility (University of California, Irvine; Irvine, CA, USA) at approximately 10 weeks of age, and began experimentation at approximately 12 weeks of age. They were pair-housed in a colony room set to a standard 12:12 hr light:dark schedule. The rats were tested during the light phase. Water was always provided ad libitum in the home cages. Rats were fed between 10-14 g of standard lab chow per day (Teklad 2020X) during the experiment to maintain them at ~85% of their estimated free-feeding bodyweight. Rats were handled for 3 days before training. Husbandry and experimental procedures were approved by the UC Irvine Institutional Animal Care and Use Committee (IACUC) and were in accordance with the National Research Council Guide for the Care and Use of Laboratory Animals.

#### 2.1.2. Apparatus

The experiment was conducted in 16 operant chambers (Med-Associates; St. Albans, VT), each housed within sound-attenuating, ventilated boxes. Each chamber was equipped with a stainless-steel grid floor; two stainless steel walls (front and back); and a transparent polycarbonate side-wall, ceiling, and door. Pellet dispensers, mounted on the outside of the operant chamber, were equipped to deliver 45-mg food pellets (Bio-Serv) to a recessed food cup centered on the lower section of the front wall. Head entries into the food receptacle were transduced by an infrared photobeam. A retractable lever was located to the left of the food cup, on the front wall. The chamber was also equipped with a house light centered at the top of the back wall. Auditory stimuli were presented to animals via a speaker located on the back wall. Experimental events were controlled and recorded with 10-ms resolution by the software program MED-PC IV (Tatham and Zurn, 1989).

#### 2.1.3. Procedure

##### Magazine training

All sessions of all phases began with the onset of the houselight. In each of two 30-minute sessions of magazine training, food pellets were delivered on a random-time (RT) 60-s schedule of food deliveries.

##### Pavlovian training

Pavlovian training involved exposure to two 10-s conditioned stimuli (CS; 3-kHz tone and 10-Hz clicker) paired with reward (grain-based food pellets). Rats were assigned to one of four groups with different CS-reward contingencies. For Group 10/90 (*n* = 8), the probability that a single food pellet would be delivered at CS offset was 10% for one cue and 90% for the other cue. For Group 30/70 (*n* = 8), the probability of reward was 30% for one cue and 70% for the other cue. For Group 50/50 (*n* = 8), the probability of reward was 50% for both cues. These arrangements allowed us to establish a range of CS-reward contingencies while controlling for the total number of rewards delivered per session. We also ran a control condition, Group 0/0 (*n* = 6), which received no reward deliveries during this phase of training, such that the probability of reward was 0% for both cues.

In each session, a 55-s interval preceded onset of the first CS. There was a fixed 85-s + variable 25-s inter-stimulus interval (ISI) between consecutive CS presentations (i.e., between previous CS offset and subsequent CS onset), and a 55-s interval following the final CS presentation prior to the end of the session. Pavlovian training lasted for 9 sessions, each involving 20 pseudorandomly-alternating presentations of each CS (40 trials total per session).

##### Instrumental training

During initial instrumental (lever-press) training, rats were continuously reinforced with a food pellet delivery for pressing the left lever (fixed-ratio, FR-1), earning a maximum of 30 pellets per session. These sessions lasted no more than 30 min. Rats were required to earn all 30 food pellets in two consecutive sessions before advancing. During subsequent training sessions, lever pressing was reinforced according to a random-interval (RI) schedule, such that the lever remained available but was inactive for an average of *t* seconds after each reward delivery, where individual t values were selected randomly from an exponential distribution. The RI schedule was changed over training days with one day of RI-5 (*t*= 5 sec), one day of RI-15 (*t* = 15 sec), two days of RI-30 (*t* = 30 sec), and six days of RI-45 (*t* = 45 sec) training. Each RI session lasted 30 minutes.

##### Pavlovian-to-instrumental transfer (PIT)

Following instrumental training, rats received one session of reminder Pavlovian training (identical to earlier sessions), and a 30-min session of instrumental extinction, in which the lever was continuously available but was inactive. Rats then received a 43-minute PIT test session, during which the lever was once again continuously available but inactive. During the test, rats received 6 noncontingent presentations of each 10-s CS in pseudorandom order (ABBABAABABBA). The ISI was 180 s, and a 6.5-min interval preceded onset of the first CS (i.e., 5 min plus one half of the ISI). No food pellets were delivered at test.

### 2.2. Experiment 2

This experiment applied a PIT protocol based on Experiment 1 to compare the behavioral responses of adolescent and adult rats when presented with cues signaling a low (30%) or high (70%) probability of reward.

#### 2.2.1. Animals and Apparatus

Thirty experimentally naïve male Long Evans rats were used in this experiment: 12 adolescents and 12 adults. Rats were derived from a local colony and weaned at PND 21-23. They were group-housed (2-3 rats per cage). Adults began testing at approximately 18 weeks of age, and adolescents began testing at PND29. Testing occurred over the span of 19 days, such that younger rats ended testing by PND47, which falls within the typical period of puberty for male rats (Schneider et al., 2008) and corresponds to a period of middle to late adolescence (Friemel et al., 2010). As in Experiment 1, the colony room was set to a standard 12:12 hr light:dark schedule, the rats were tested during the light phase, and water was always provided ad libitum in the home cages. Food was also provided ab libitum up until two days before the beginning of the experiment, after which rats were provided lab chow to maintain at them at ~85% of free-feeding body weight, corrected for growth. The experiment was conducted in the same chambers and using the same materials as Experiment 1, except that sucrose pellets were used as the reinforcer. Rats were handled for 3 days before training and were given 1 day of pre-training exposure to sucrose pellets to attenuate neophobia. Husbandry and experimental procedures were approved by the UC Irvine Institutional Animal Care and Use Committee (IACUC) and were in accordance with the National Research Council Guide for the Care and Use of Laboratory Animals.

#### 2.2.2. Procedure

Experiment 2 was similar to Experiment 1, with the following exceptions identified below. Generally, the entire procedure was abbreviated relative to Experiment 1 to ensure that behavioral testing was restricted to the peripubertal period. Accordingly, for instance, instrumental training ended with reinforcement on an RI-30 schedule. Further, as described above, we used sucrose (not grain-based) pellets as food reward in Experiment 2. We switched to this more palatable reward to facilitate task acquisition, particularly given that we used a modest food deprivation regimen to allow for normal growth in developing rats. We also wished to facilitate comparisons with our past research on the changes in sucrose consummatory behavior during adolescence (Marshall et al., 2017b).

##### Magazine training

Magazine training was identical to Experiment 1, except that it lasted for only one session.

##### Pavlovian training

Pavlovian training was identical to Experiment 1, except that all rats were exposed to the conditions of Group 30/70 in Experiment 1. Additionally, Pavlovian training lasted for only 7 days.

##### Instrumental training

Instrumental training was identical to Experiment 1, with the following exceptions. FR1 training ended when each rat had earned 30 pellets within one session. Nine adolescent rats and 10 adult rats achieved this criterion within one session; 2 adolescent rats required 2 sessions, 1 adolescent rat required 3 sessions, 1 adult rat required 4 sessions, and 1 adult rat required 8 sessions. Subsequently, rats were given one day of RI-5 training, one day of RI-15 training, and 5 days of RI-30 training. The adult rat who required 8 FR-1 training sessions was given only 4 sessions of RI-30 training to ensure that all rats were tested together (on PND47 in the adolescent group).

##### PIT

Rats received one session of reminder Pavlovian training and a 30-min session of instrumental extinction, which was followed by a PIT test session. These procedures were identical to those described in Experiment 1.

### 2.3. Data Analysis

All summary measures were obtained from the raw data using MATLAB (The MathWorks; Natick, MA, USA), and analyzed with mixed-effects regression models (Pinheiro and Bates, 2000), a powerful analytical framework that is both well established and highly recommended for behavioral research (Boisgontier and Cheval, 2016). Mixed-effects models are comparable to repeated-measures regression analyses, and allow for parameter estimation per manipulation condition (fixed effects) and the individual (random effects) (Bolker et al., 2009; Hoffman, 2013; Hoffman and Rovine, 2007; Pinheiro and Bates, 2000; Schielzeth and Nakagawa, 2013). Mixed-effects regression models (1) effectively handle missing data and (2) permit the inclusion of categorical and continuous predictors in the same analysis, thus allowing detection of group-level changes across ordered data samples (i.e., continuous time points) while also accounting for corresponding individual differences. All relevant fixed-effects factors were included in each model, and model selection of random-effects terms was performed using the Akaike information criterion (AIC), in which the doubled negative log likelihood of the model is penalized by twice the number of estimated parameters (Burnham and Anderson, 2002). Categorical predictors were effects-coded (i.e., codes sum to 0), and continuous predictors were mean-centered (Kreft et al., 1995). For Experiment 1, the fixed-effects structure of the analyses of Pavlovian training and PIT included main effects of group and reward probability; for Experiment 2, the corresponding fixed-effects structure included the main effects of group and reward probability as well as the group-by-reward probability interaction. Instrumental training analyses incorporated generalized linear mixed-effects models (family: gamma, link: log) with predictors of group and time since the previous reward delivery. The alpha level for all tests was .05.

Effect size was represented by the unstandardized regression coefficient (Baguley, 2009), reported as *b* in model output tables. The source of significant interactions was determined by post hoc marginal *F* tests using MATLAB’s *coefTest* function. Main effects and interactions are reported in-text as the results of ANOVA *F*-tests (i.e., whether the coefficients for each fixed effect were significantly different from 0). Full model output and specification of random-effects structures are provided in *Supplemental Information*.

Our primary dependent measures were lever pressing and food-cup approach behavior. Because the behavioral effects of reward-paired cues often persist into the post-cue period (Delamater and Holland, 2008; P F Lovibond, 1983; Marshall and Ostlund, 2018), we quantified cue-induced changes in behavior by subtracting the mean response rate during local pre-CS periods (10 sec each) from the mean response rate during 20-sec periods beginning at CS onset and extending 10 sec after CS offset. Pre-CS (baseline) data were averaged across all CS trials (within subject). We also calculated a *response bias* measure to quantify how CS presentations altered the way rats distributed their activity between these two responses. Specifically, cue-elicited food-cup approach rate (CS – pre-CS) was subtracted from cue-elicited press rate (CS – pre-CS), such that positive values indicated a bias toward the food cup and negative values indicated a bias toward the lever. Importantly, food-cup approach behavior can fall into two categories: spontaneous approaches and approaches that are performed as part of an instrumental *press-approach* action sequence (Halbout et al., 2019; Marshall and Ostlund, 2018), the latter indicated by increased likelihood of food-cup approach shortly after lever presses (Supplemental Figures 1-2). Accordingly, as done previously, we excluded approaches that occurred within a 2.5 sec post-lever press period from our analysis.

When necessary, dependent variables were square-root transformed to correct for positive skewness. If square-root transformations were unable to adequately correct for skewness, the data were Yeo-Johnson transformed (Yeo and Johnson, 2000) using the *bestNormalize* package in R (Peterson and Cavanaugh, 2019). Data points of difference scores were removed if their values were at least three scaled median absolute deviations from the median (Leys et al., 2013). Notably, because we used regression analyses, data point removal due to outlier identification did not require the animal to be removed from analysis, just the outlying data point. Further, this only occurred in Experiment 2: two data points for Pavlovian training analysis (one adult 30% CS, one adult 70% CS), two data points for PIT lever press analysis (two adult 70% CS), two data points for PIT food-cup approach analysis (one adult 30% CS, one adult 70% CS), and two data points for PIT response bias analysis (two adult 70% CS).

Figures incorporated nontransformed data points for ease of interpretation; transformed data, along with individual rats’ data points, are provided in *Supplemental Information*. For both experiments, the final three sessions of Pavlovian training were used to assess conditioned approach behavior during CS+ and CS− trials relative to pre-CS baseline periods. Analyses of instrumental training included the final three sessions of training.

## 3. Results

### 3.1. Experiment 1

We conducted an initial PIT experiment to determine the influence of expected reward probability on the way that normal adult male rats distribute their activity between lever pressing and food-cup approach behavior. Our goal was to identify cue conditions that evoke distinct response tendencies and would therefore be useful for probing cue-elicited lever pressing across varying levels of response competition with food-cup approach behavior. Based on our recent research (Marshall et al., 2018) and related findings (see Discussion), we predicted that cues signaling a low probability of reward would be most effective in eliciting lever pressing and least effective in eliciting food-cup approach. In contrast, cues associated with a high probability of reward were expected to interfere with this impulse to lever press and instead elicit food-cup approach behavior. Our plan was to use the findings of this experiment to develop a PIT protocol for characterizing behavioral differences between adolescent and adult rats in Experiment 2.

#### 3.1.1. Pavlovian and instrumental training

During Pavlovian conditioning, the probability that a CS would co-terminate with the delivery of a food pellet varied across cues and groups. Figure 1A shows the mean CS-induced increase in food-cup approach behavior during the final 3 Pavlovian training sessions. Group 0/0, for which neither cue signaled food delivery, showed essentially no CS-induced approach behavior (data collapsed across CSs). Similarly, Group 10/90 did not increase their food-cup approach behavior when presented with the CS signaling a 10% reward probability. Aside from these conditions, all other CSs elicited an increase in food-cup approach. Linear mixed-effects analysis of these difference scores (square-root transformed) confirmed our impression that CS-elicited approach responses increased with expected reward probability, *F*(1, 55) = 48.52, *p* < .001 (Supplemental Table 1, Supplemental Figure 3).

**Figure 1.**
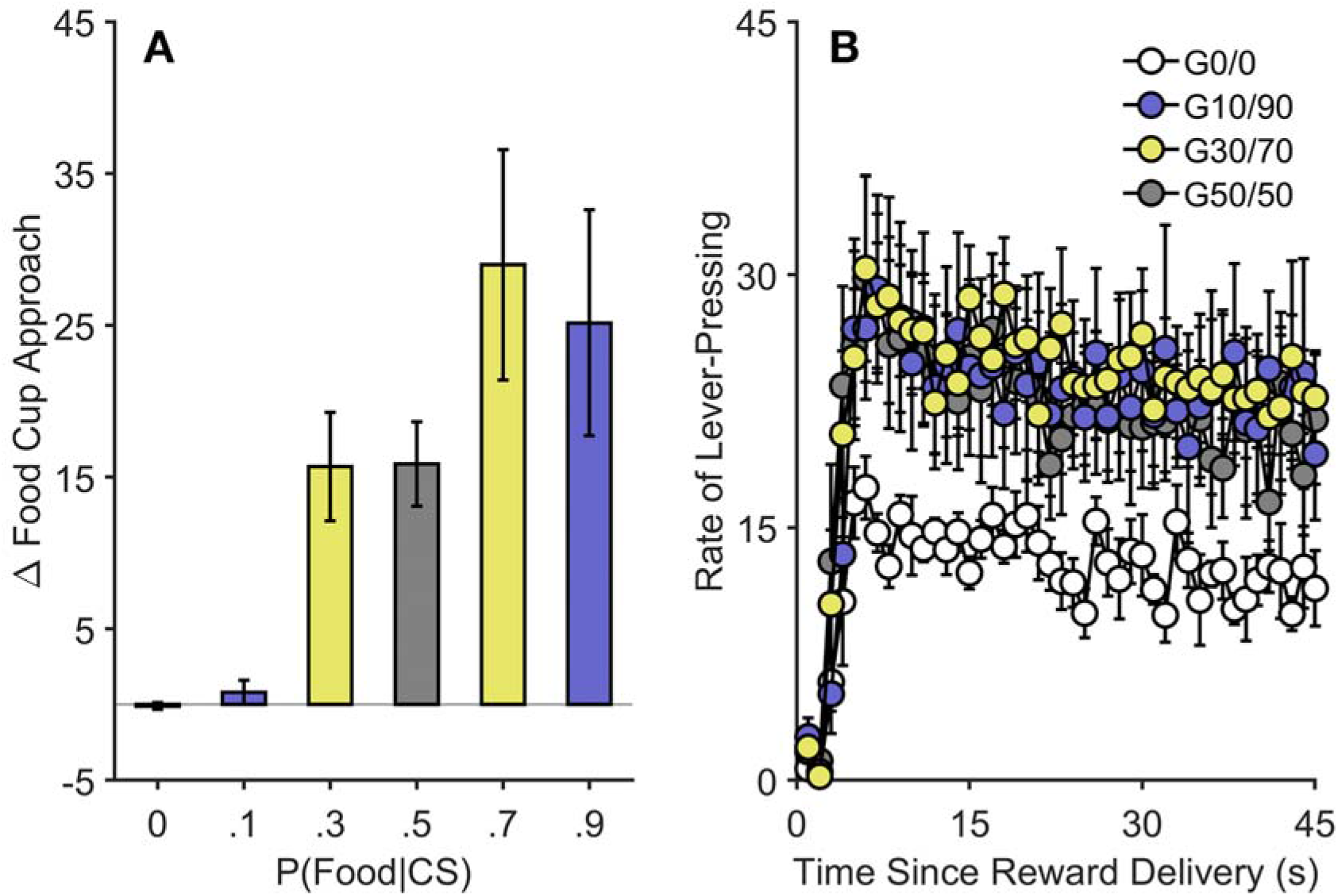
Pavlovian and instrumental training in Experiment 1. (A) Cue-induced changes in food-cup approach behavior increased with expected reward probability. Data are averaged over the final 3 days of Pavlovian training. Data represent the rate of food cup approaches (response per minute) during the pre-cue period subtracted from the rate of food cup approaches during the cue period. (B) Response gradients during the final 3 days of instrumental training. Except for Group 0/0 (G0/0), all groups responded at similar rates. Data represent the rate of lever pressing (i.e., responses per minute, controlling for the number of opportunities to respond in each time bin). See Methods for details. Food cup approach difference scores were square-root transformed for analysis but plotted in nontransformed space for ease of interpretation. Error bars reflect ± 1 between-subjects standard error of the mean. CS = conditioned stimulus. G = group. P = probability.

Following Pavlovian training, rats were trained to perform an instrumental lever-press response for food reward, which was ultimately reinforced according to an RI 45-s schedule, such that rats had to wait an average of 45-sec after each reinforced lever press before the next reinforcer could be earned. Figure 1B shows mean lever-press rates for each group as a function of time since the previous reward delivery, collapsed across the final 3 sessions of RI-45 training. Interestingly, even though all rats responded for the same reward on the same schedule, Group 0/0’s asymptotic rate of lever pressing was considerably lower than that of the other groups. For statistical analysis, we removed the first 10 s of data after reward delivery to allow response rates to restabilize after reward consumption. In line with our initial impression, the best-fitting generalized linear mixed-effects model revealed a main effect of group, *F*(3, 1045) = 3.05, *p* = .028, and post-hoc marginal *F* tests indicated that Group 0/0 responded at a significantly lower rate compared to the other three groups, *p*s ≤ .030, and that Groups 50/50, 30/70, and 10/90 did not significantly differ, *p*s ≥ .456 (Supplemental Table 2, Supplemental Figure 4).

#### 3.1.2. PIT

We then conducted a PIT test to probe the influence of reward-predictive cues on rats’ tendency to perform the lever-press and food-cup approach responses and determine whether this influence varies as a function of expected reward probability. During PIT tests, rats had continuous access to the lever and food cup alcove but received no reward deliveries, which led to gradual extinction of both responses over time. During the session, each 10-s CS was noncontingently presented to determine its impact on behavior relative to baseline.

Figure 2 presents the results of PIT testing. As can be seen in Fig 2A, the effect of CS presentations on lever-press performance varied as a function of expected reward probability, *F*(1, 55) = 17.04, *p* < .001 (Supplemental Table 3, Supplementals Figure 5-6). Post hoc marginal *F* tests indicated that there was no significant change in lever pressing (relative to pre-CS period) following the 0%, 50%, 70%, or 90% CSs, *p*s ≥ .152. For Group 10/90, the 10% CS elicited a significant increase in lever pressing, *p* = .004, and a similar trend was found for the 30% CS in Group 30/70, *p* = .053. Cues that signaled a low but non-zero probability of reward were therefore most effective in eliciting an increase in lever-press performance. In contrast, as shown in Fig 2B, cue-induced changes in food-cup approach (square-root transformed) increased as a function of expected reward probability, *F*(1, 55) = 63.99, *p* < .001 (Supplemental Table 4, Supplemental Figures 5-6). Post hoc marginal *F* tests indicated that there was no significant cue-elicited change in food-cup approach following the 0% and 10% CSs, *p*s ≥ .737. However, there were significant increases in cue-elicited food-cup approach during the 30%, 50%, 70%, and 90% CSs (*p*s < .001).

**Figure 2.**
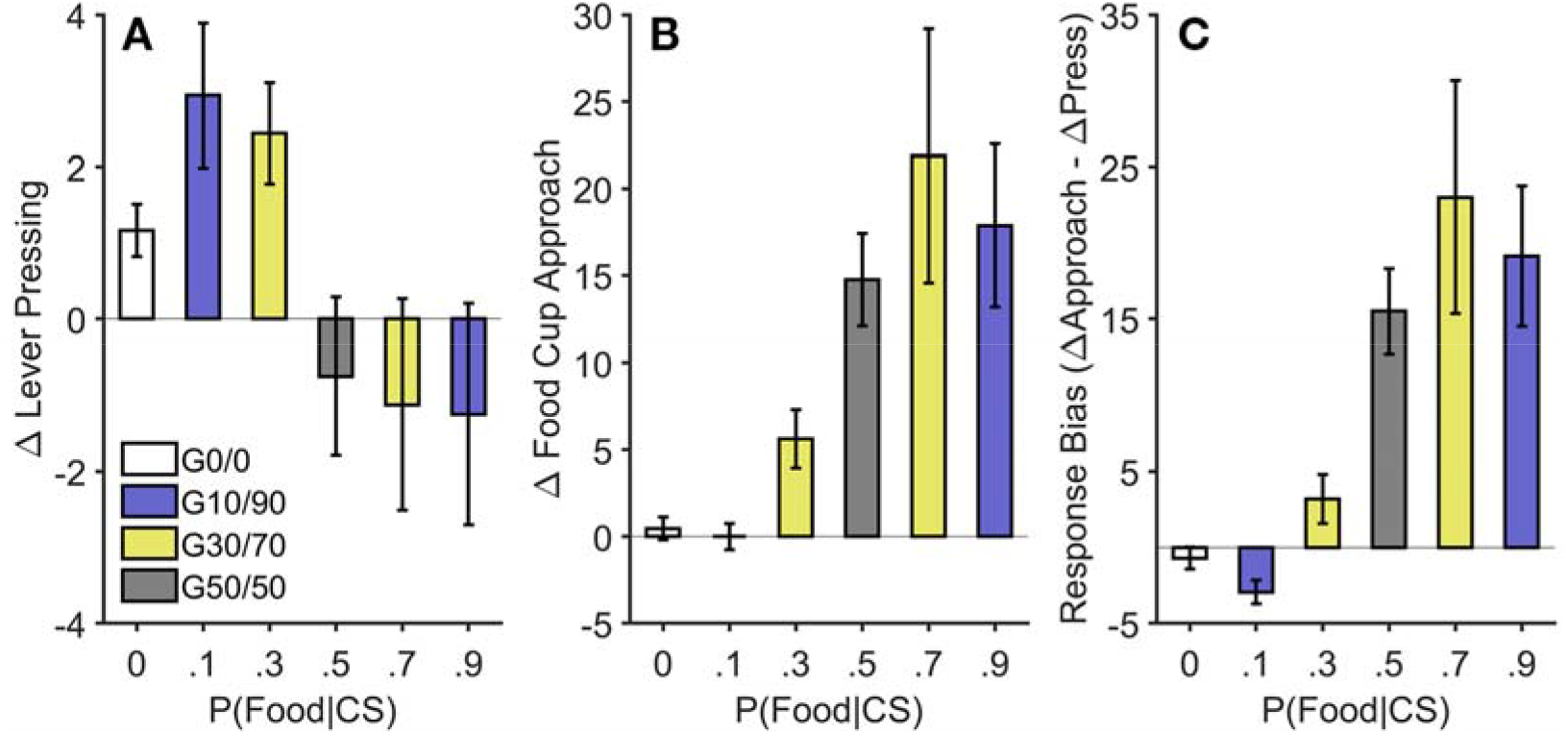
Pavlovian-to-instrumental transfer in Experiment 1. (A) Cues that signaled a low probability of reward were more effective at eliciting lever pressing than cues signaling a high probability of reward. Data represent the rate of lever pressing (i.e., responses per minute) during the pre-cue period subtracted from the rate of lever pressing during the cue period and 10-s post-cue period. (B) Concurrent changes in food-cup approach behavior during CS presentations increased with expected reward probability. Data represent the rate of food cup approach (i.e., responses per minute) during the pre-cue period subtracted from the rate of food cup approach during the cue period and 10-s post-cue period. (C) The tendency for cues to bias behavior toward the food cup relative to the lever increased with expected reward probability.

Lastly, we contrasted the effects of the cues on food-cup approach and lever pressing using a *response bias* score (CS-induced approach – CS-induced pressing), which is presented in Figure 2C. Mixed model analysis of these data (Yeo-Johnson transformed to correct for positive skewness) found that rats’ bias toward the food cup increased with the strength of the CS-reward probability, *F*(1, 55) = 101.54, *p* < .001 (Supplemental Table 5). Thus, when presented with a CS that signaled a high probability of reward, rats withheld their lever-press performance and instead focused their behavior at the food cup.

Data represent cue-induced changes in lever pressing (A) subtracted from cue-induced changes in food cup approach (B). See Methods for details. For analyses, food cup approach difference scores (B) were square-root transformed for analysis, and response bias data (C) were Yeo-Johnson transformed. Both are plotted in nontransformed space for ease of interpretation. Error bars reflect ± 1 between-subjects standard error of the mean. CS = conditioned stimulus. G = group. P = probability.

### 3.2. Experiment 2

Having established effective conditions for contrasting the distinct behavioral effects of weak versus strong reward-predictive cues, we next investigated if adolescent and adult rats differed in the way they responded to such cues. Our PIT protocol was based on Group 30/70 from Experiment 1, which showed a clear cue-specific pattern of responding, increasing their lever-press performance during the low-probability cue and withholding this response in order to check the food cup during the high-probability cue. This approach allowed us to efficiently assess the motivational influence of reward-paired cues under conditions with (70% CS) and without (30% CS) a strong competing response tendency.

#### 3.2.1. Pavlovian and instrumental training

Adolescent (*n* = 12) and adult rats (*n* = 12) were given differential Pavlovian training with cues signaling either a low (30%) or high (70%) probability of reward. Figure 3A shows the mean rate of cue-elicited food-cup approach during the final three days of training. Two data points were identified as outliers and removed from the analysis. As in Experiment 1, linear mixed model analysis found a significant effect of expected reward probability, *F*(1, 42) = 13.93, *p* = .001, indicating that the 70% CS was generally more effective at eliciting approach (square-root transformed) than the 30% CS (Supplemental Table 6, Supplemental Figure 7). The influence of expected reward probability was not moderated by group, *F*(1, 42) = 2.21, *p* = .145, nor was there a main effect of group, *F*(1, 42) = 0.82, *p* = .370.

**Figure 3.**
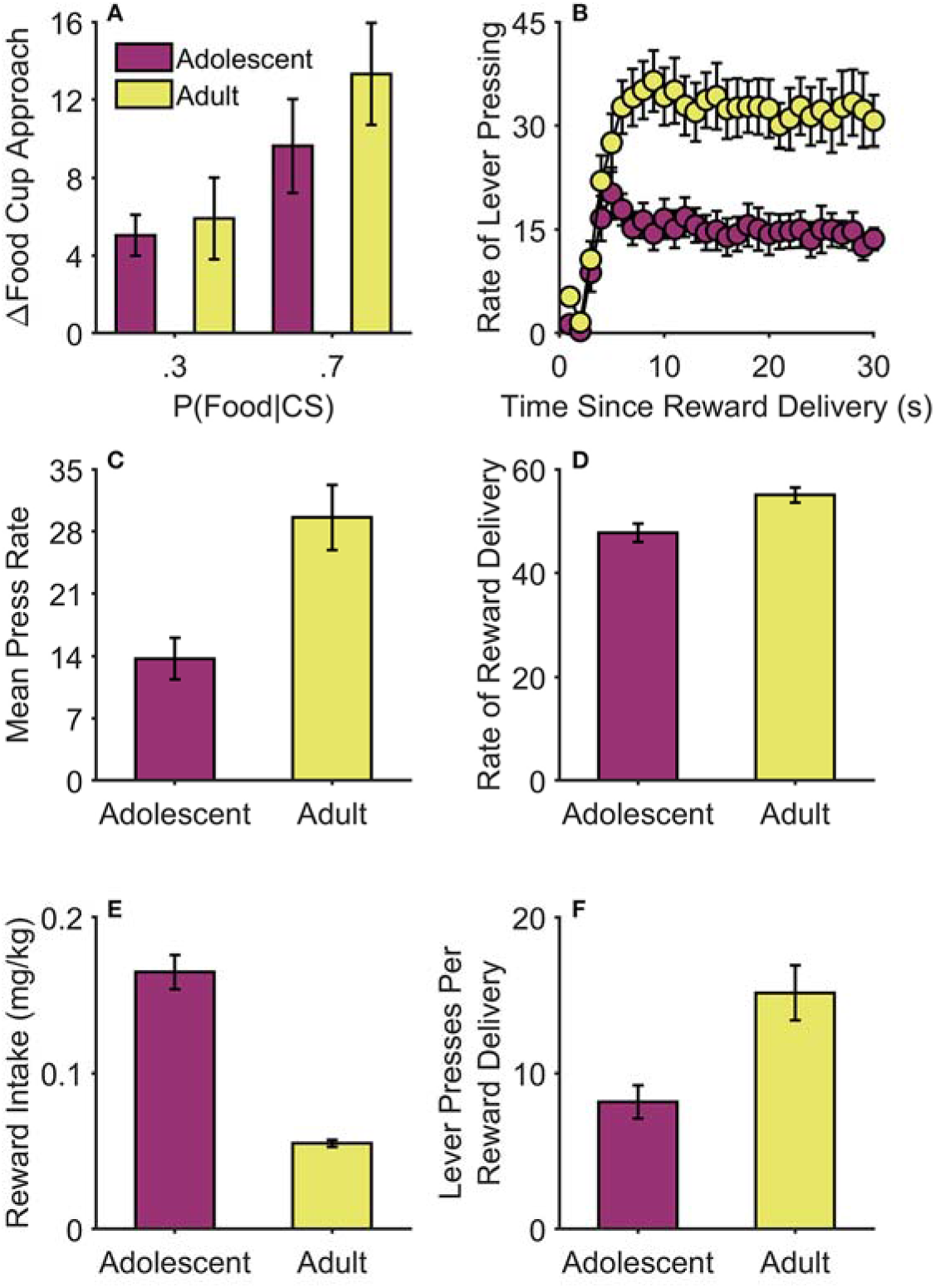
Pavlovian and instrumental training in Experiment 2. (A) Cue-induced changes in food-cup approach behavior increased with expected reward probability in both adults and adolescents. Data are from the final 3 days of Pavlovian training and represent the rate of food cup approaches (response per minute) during the pre-cue period subtracted from the rate of food cup approaches during the cue period. (B) Response gradients during the final 3 days of instrumental training. Data represent the rate of lever pressing (i.e., responses per minute, controlling for the number of opportunities to respond in each time bin). Overall, adolescents responded at lower rates than adults (C). Adults experienced a greater rate of reward delivery than adolescents (D); data represent mean number of rewards earned per session. In contrast, weight-adjusted reward intake (mg of food pellets per kg of body weight) per session was considerably higher in adolescents (E). Adolescents were more efficient in their responding than adults, exhibiting fewer lever presses per reward delivery (F). See Methods for details. Food cup approach difference scores were square-root transformed for analysis but plotted in nontransformed space for ease of interpretation. Error bars reflect ± 1 between-subjects standard error of the mean. CS = conditioned stimulus. P = probability.

In contrast, the mean rate of lever pressing during the last three days of instrumental training was significantly lower in the adolescent group relative to adults, which was apparent when the analysis excluded data from the 10-s post-reinforcement period (Figure 3B, *F*(1, 476) = 17.86, *p* < .001; Supplemental Table 7, Supplemental Figure 7), or was averaged over the entire interval (Figure 3C, *z* = 3.15, *p* = .002 (Wilcoxon rank-sum test). Adults also earned more food pellets per session than adolescents (Figure 3D), *z* = 2.72, *p* = .007 (Wilcoxon rank-sum test). However, weight-adjusted reward intake (i.e., mg of food per kg of body weight; Figure 3E) was significantly elevated for the adolescent group, *z* = 4.13, *p* < .001 (Wilcoxon rank-sum test). It is also worth noting that because the RI-45 schedule required only a relatively low rate of responding to maximize reward, adolescent rats were able to earn rewards more efficiently, performing fewer presses per reward delivery than adults (Figure 3F), *z* = 2.86, *p* = .004 (Wilcoxon rank-sum test) (Supplemental Figure 7).

#### 3.2.2. PIT

Adolescent and adult rats were then administered a PIT test to assess how the reward-predictive cues influence their lever-press and food-cup approach behavior. Figure 4A shows the effects of CS presentations on lever-press performance. Linear mixed-effects model analysis revealed a main effect of group, *F*(1, 42) = 4.48, *p* = .040, indicating that cue-elicited lever pressing was generally elevated in adolescent rats relative to adults (Supplemental Table 8, Supplemental Figures 8-9). While the main effect of expected reward probability (CS-type) did not reach significance, *F*(1, 42) = 2.99, *p* = .091, the influence of this factor differed between the two groups (CS-type × Group interaction), *F*(1, 42) = 4.17, *p* = .047. Post hoc marginal *F* tests found that adult rats displayed lower levels of pressing on trials with the 70% CS than the 30% CS, *p* = .014, much like adult rats in Experiment 1. In contrast, expected reward probability did not significantly influence cue-elicited lever pressing in the adolescent group, *p* = .818. The groups did not significantly differ in their baseline (pre-CS) rates of lever pressing (Adolescents: mean = 4.17, SEM = 0.66, Adults: mean = 5.52, SEM = 1.07), *z* = 1.09, *p* = .275 (Wilcoxon rank-sum test).

**Figure 4.**
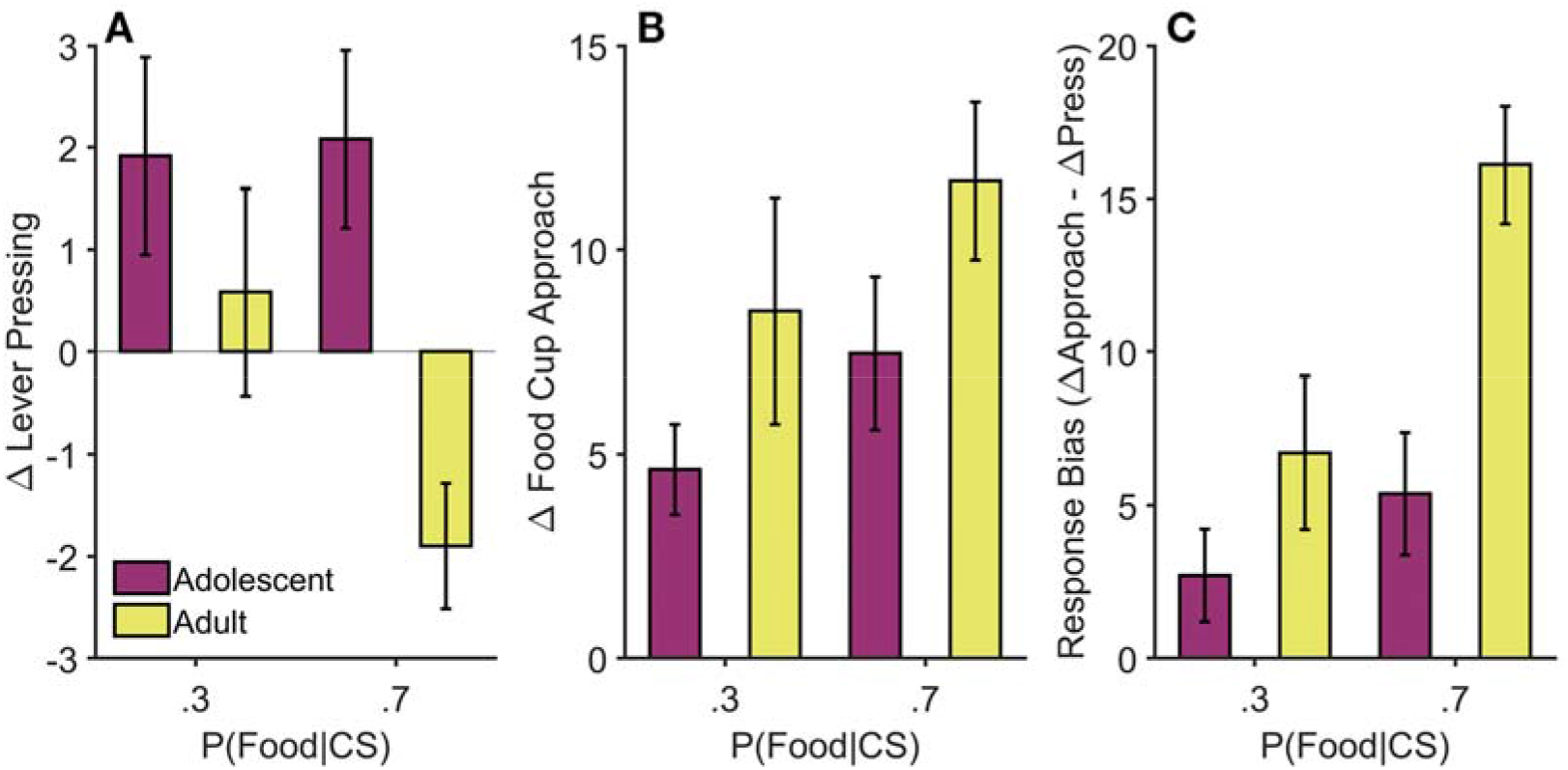
Pavlovian-to-instrumental transfer in Experiment 2. (A) For adult rats, cue-elicited lever pressing was greater during trials with the 30% CS than the 70% CS. In contrast, adolescent rats showed a similar increase in lever pressing to both cues. Data represent the rate of lever pressing (i.e., responses per minute) during the pre-cue period subtracted from the rate of lever pressing during the cue period and 10-s post-cue period. (B) Both groups showed similar patterns of cue-elicited food-cup approach behavior, though adolescent rats showed a marginally lower rate of conditioned food-cup approach. Data represent the rate of food cup approach (i.e., responses per minute) during the pre-cue period subtracted from the rate of food cup approach during the cue period and 10-s post-cue period. (C) The tendency for cues to bias behavior toward the food cup relative to the lever was greater for adults than for adolescents, particularly during the 70% CS. Data represent cue-induced changes in lever pressing (A) subtracted from cue-induced changes in food cup approach (B). See Methods for details. For analyses, food cup approach difference scores (B) and response bias data (C) were Yeo-Johnson transformed. Both are plotted in nontransformed space for ease of interpretation. Error bars reflect ± 1 between-subjects standard error of the mean. CS = conditioned stimulus. P = probability.

Figure 4B shows cue-elicited food-cup approach during PIT testing. Mixed model analysis of these data (Yeo-Johnson transformed) found trends toward main effects of group, *F*(1, 42) = 3.95, *p* = .053, and CS reward probability, *F*(1, 42) = 3.12, *p* = .085, with no significant interaction between these factors, *F*(1, 42) = 0.07, *p* = .794 (Supplemental Table 9, Supplemental Figures 8-9). The groups did not significantly differ in their rate of food-cup approach during pre-CS periods (Adolescents: mean = 2.38, SEM = 0.34, Adults: mean = 4.27, SEM = 1.17), *z* = 0.34, *p* = .731 (Wilcoxon rank-sum test).

The response bias measure (Figure 4C) shows more directly that adolescent rats differed from adults in the way they changed their activity between the lever and food cup in response to reward-predictive cues. Mixed model analysis of these data (Yeo-Johnson transformed) revealed a significant main effect of expected reward probability, *F*(1, 42) = 11.91, *p* = .001, with the 70% CS eliciting a stronger shift in responding toward the food cup than the 30% CS (Supplemental Table 10, Supplemental Figure 8). The general shift toward the food cup was significantly weaker in adolescent compared to adult rats (main effect of age group), *F*(1, 42) = 11.09, *p* = .002. While the Group × CS Type interaction did not reach significance, *F*(1, 42) = 3.56, *p* = .066, inspection of the data suggests that adolescent rats were less likely to adjust their response allocation based on cue-evoked reward predictions. Post hoc *F* tests confirmed that for adult rats the 70% CS elicited a stronger bias toward the food cup than the 30% CS, *p* < .001, whereas adolescent rats did not significantly adjust their response bias based on expected reward probability, *p* = .261.

## 4. General Discussion

The current study investigated the behavioral underpinnings of adolescent impulsive behavior using a new PIT protocol designed to probe impulse control over cue-motivated reward seeking in rats. We show that the tendency for a reward-paired cue to motivate exploratory reward-seeking behavior (instrumental lever pressing) is strongly modulated by its predictive value. Adult rats increased their rate of lever pressing when presented with a cue that signaled a low probability of reward but withheld this behavior and instead approached the food cup when presented with a more predictive cue. In contrast, adolescent rats were impaired in using expected reward probability to modulate their reward-seeking behavior, increasing their rate of lever pressing in response to both weak and strong reward predictors. As discussed below, we suggest that this heightened motivational response to reward-paired cues in adolescent rats is driven by an imbalance between impulse control and motivational systems.

Researchers have long recognized that the predictive value of reward-paired cues determines the type of the conditioned responses that they come to elicit, with weak predictors stimulating general foraging behaviors and strong predictors eliciting a more narrow set of responses required for retrieving and consuming the expected reward (Bindra, 1974; Konorski, 1967; Timberlake et al., 1982). The current findings are in line with this view and with previous PIT studies showing that instrumental reward-seeking behavior is facilitated by weak cues (Estes, 1948, 1943; Meltzer and Brahlek, 1970) but is suppressed by cues that signal imminent reward (Azrin and Hake, 1969; Lovibond, 1981; Van Dyne, 1971). However, strong predictors of food reward have been shown to acquire latent motivational properties, which can be unmasked by treatments that weaken the expression of the competing food-cup approach response (Baxter and Zamble, 1982; Holmes et al., 2010; Peter F. Lovibond, 1983). Withholding general foraging behavior when there is a strong reward expectancy is adaptive because it helps conserve energy and minimize the risk of reward loss (see Mackintosh, 1974; Timberlake et al., 1982). This ability to flexibly suppress motivational impulses when they become maladaptive is a defining feature of what some have termed ‘hot’ or affective cognitive control (Metcalfe and Mischel, 1999; Ochsner and Gross, 2005). We propose that both weak and strong reward-predictive cues have the capacity to motivate general reward seeking, but that strong predictors are unique in that they also inhibit this impulse to allow for efficient retrieval of the expected reward.

Further research is needed to more fully characterize the nature of response conflict between lever pressing and food-cup approach behavior during PIT testing. We propose that the PIT paradigm can serve as a *naturalistic go/no-go task*, in which weak cues motivate lever pressing and strong cues actively *inhibit* this impulse in favor of checking the food cup. However, it remains to be determined whether competition between these behaviors is resolved through top-down inhibition of cue-motivated behavior or some other method of arbitration between systems controlling instrumental and Pavlovian behavior (Dayan et al., 2006; Dorfman and Gershman, 2019; Guitart-Masip et al., 2012). This may depend on the specific behavioral strategies that rats use when lever pressing and checking the food cup. The random interval training protocol used for instrumental training here and in similar PIT studies is known to promote to the development of instrumental reward-seeking habits, which are automatically performed without considering the value of anticipated outcomes (de Russo et al., 2010; Dickinson, 1985; Yin et al., 2004). Other findings indicate that such habits are more sensitive than goal-directed actions to the motivational influence of reward-paired cues, as measured by the PIT effect, and that this influence is, itself, insensitive to devaluation (Colwill and Rescorla, 1988; Holland, 2004; Rescorla, 1994; Wiltgen et al., 2012). In contrast, reward devaluation studies indicate that conditioned food-cup approach behavior is not performed habitually (or reflexively), but instead depends on cue-evoked reward expectations, even after extensive Pavlovian training (Holland, 1998; Holland and Rescorla, 1975; Keefer et al., 2020). Although we did not confirm these differential behavioral effects of reward devaluation in the current study, such findings suggests that strong reward predictors engage cognitive processes that are able override the implicit incentive motivational processes that underlie the PIT effect.

Cognitive control is believed to be particularly important when complex situational cues must be used to resolve conflict between competing response tendencies (Miller, 2000). This aspect of cognitive control has recently been linked to the tendency for cues to bias action selection based on sensory-specific features of the expected reward (Balleine, 2016), a phenomenon referred to *as outcome-specific PIT*. We propose that cognitive control processes may also guide action selection in PIT based on other information encoded about the reward. The current study provides a parametric demonstration that expected reward probability influences the degree to which rats choose to lever press versus check the food cup. Similarly, we recently found that rats flexibly adjust their response preferences over time during cues that signal delayed reward, transitioning from the lever to the food cup as the expected reward delivery time approaches (Marshall and Ostlund, 2018).

Based on this theoretical framework, adult rats in the current study displayed good cognitive control, flexibly adjusting their response bias to promote lever pressing when the expected probability of reward was low (30% CS trials) and food-cup approach when it was high (70% CS trials). In contrast, adolescent rats showed a diminished capacity for cognitive control, increasing their rate of lever pressing even when presented with the high-probability cue, which interfered with their ability to perform the more adaptive food-cup approach response. If adolescent rats were simply more motivated by reward-paired cues (with a normal capacity for cognitive control), then they should have exhibited higher levels of cue-elicited lever pressing when this was adaptive (30% CS trials), but should have retained an ability to inhibit lever pressing when it was maladaptive (70% CS trials). This might explain why previous studies have generally not found evidence of elevated cue-motivated behavior in adolescent rats on tasks that do not involve an explicit cognitive control requirement (Anderson et al., 2013; Anderson and Spear, 2011; Doremus-Fitzwater and Spear, 2011; Naneix et al., 2012).

Our findings illustrate the central role that cognitive control dysfunction plays in adolescent impulsive behavior and provides an approach for modeling this phenomenon. However, it remains possible that developmental changes in motivation also contributed to this effect. For instance, cues may have triggered exceptionally strong motivation to pursue reward in adolescent rats, which may have overwhelmed their limited capacity to withhold exploratory reward seeking. This possibility is in line with theories that link adolescent impulsive behavior to a developmental imbalance between transiently hyperactive emotional and motivational systems and still maturing cognitive control systems (Casey et al., 2008; Ernst et al., 2006; Steinberg, 2004). Indeed, research in humans has shown that adolescents are particularly prone to risk taking when tested in “hot” affective contexts designed to elicit arousal as opposed to “cold” deliberative contexts (Cauffman et al., 2010; Figner et al., 2009). This account is also in line with animal studies showing that adolescent rats display heightened emotional and motivational responses to palatable food rewards (Friemel et al., 2010; Marshall et al., 2017b; Schneider et al., 2015; Stolyarova and Izquierdo, 2015; Wilmouth and Spear, 2009). We suggest that this upregulation in reward processing likely exacerbates cognitive control impairment in adolescent rats, leading to maladaptive cue-motivated behavior described here.

The response conflict PIT protocol used here may prove useful in elucidating neural mechanisms of normal and impaired cognitive control. A large and growing body of work using conventional PIT protocols has identified many elements of the neural circuitry mediating the motivational influence of reward-paired cues, which includes the mesolimbic dopamine system, the nucleus accumbens, and the central amygdala (Cartoni et al., 2016; Corbit and Balleine, 2016). Research on impulse control suggests that frontal areas such as the medial prefrontal cortex and anterior cingulate provide top-down regulation over this circuitry to suppress maladaptive reward-seeking actions (Andrzejewski et al., 2011; Dalley et al., 2011). While disrupting medial prefrontal cortex function does not alter PIT expression when lever pressing is motivated by relatively weak reward-predictive cues (Cardinal et al., 2003; Corbit and Balleine, 2003; Halbout et al., 2019), it remains to be determined if this region may be preferentially engaged by strong reward predictors in order to suppress ongoing lever pressing and facilitate food-cup approach. Given that the prefrontal cortex does not establish mature patterns of connectivity until early adulthood (Andersen et al., 2000; Cressman et al., 2010; Delevich et al., 2018), future research should investigate if the heightened cue-elicited reward-seeking behavior displayed by adolescent rats in the current study reflects a failure to fully engage these mechanisms of top-down cognitive control.

The current findings are interesting to consider in relation to previous reports that repeated exposure to psychostimulant drugs can potentiate expression of PIT (LeBlanc et al., 2014, 2013; Ostlund et al., 2014; Saddoris et al., 2011; Wyvell and Berridge, 2001). This drug-induced augmentation of cue-elicited reward seeking is associated with altered task-related neural activity and phasic dopamine signaling in the nucleus accumbens (Ostlund et al., 2014; Saddoris et al., 2011). However, as we have previously noted (Marshall and Ostlund, 2018), much of this research has been conducted using modified PIT procedures involving strong cues that signal imminent reward. Under these conditions, we find that rats with a history of cocaine exposure exhibit a maladaptive increase in lever pressing and a reduction in food cup approach behavior, relative to drug-naïve rats (Marshall and Ostlund, 2018). This behavioral profile strongly resembles the altered response bias displayed by adolescent rats in the current study and is indicative of a deficit in cognitive control, perhaps in addition to a more general upregulation in incentive motivation.

Whether resulting from adolescent developmental changes or drug exposure, the inability to regulate cue-motivated behavior may promote impulsive reward seeking, potentially creating a vicious cycle when it leads to adolescent drug use. This possibility is supported by both human and non-human animal research suggesting that adolescent drug use may stimulate the development of drug addiction (Anker and Carroll, 2010; Chambers et al., 2003; Doremus-Fitzwater et al., 2010; Kandel et al., 1992; Reboussin and Anthony, 2006; Wong et al., 2013). The current findings may help guide future research investigating this problem and its underlying neurobiology.

## Acknowledgments

Conceived and designed the experiments: ATM, NTM, and SBO. Performed the experiments: ATM. Analyzed the data: ATM and SBO. Wrote the paper: ATM, NTM, and SBO.

