## Supplemental Figures for "Reward-predictive cues elicit maladaptive reward seeking in adolescent rats"

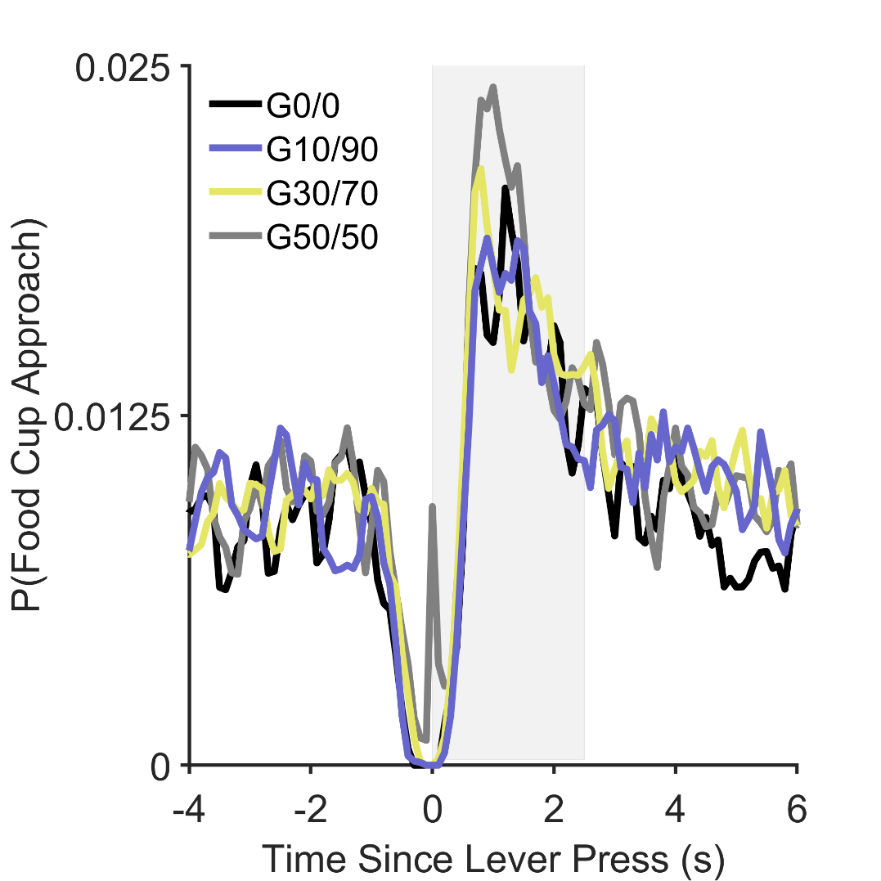

**Supplemental Figure 1**. Probability of food cup approach as a function of time from each lever press in Experiment 1. Data are from the instrumental extinction session preceding the Pavlovian-to-instrumental transfer (PIT) test. The shaded area represents 2.5 s since a lever press, the interval during which subsequent food cup approaches were designated as being part of the reward-seeking – food-cup-approach sequence. These food-cup approaches were not included in analysis of cue-induced conditioned food-cup approach within the PIT test.

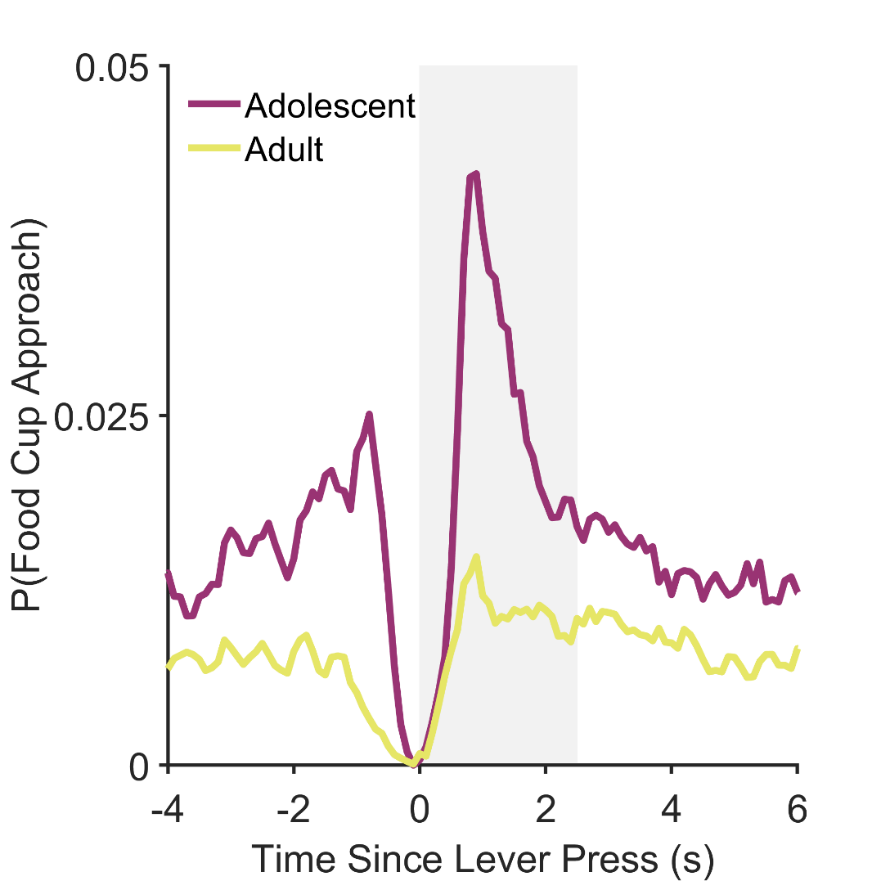

**Supplemental Figure 2**. Probability of food cup approach as a function of time from each lever press in Experiment 2. Data are from the instrumental extinction session preceding the Pavlovian-to-instrumental transfer (PIT) test. The shaded area represents 2.5 s since a lever press, the interval during which subsequent food cup approaches were designated as being part of the reward-seeking – food-cup-approach sequence. These food-cup approaches were not included in analysis of cue-induced conditioned food-cup approach within the PIT test.

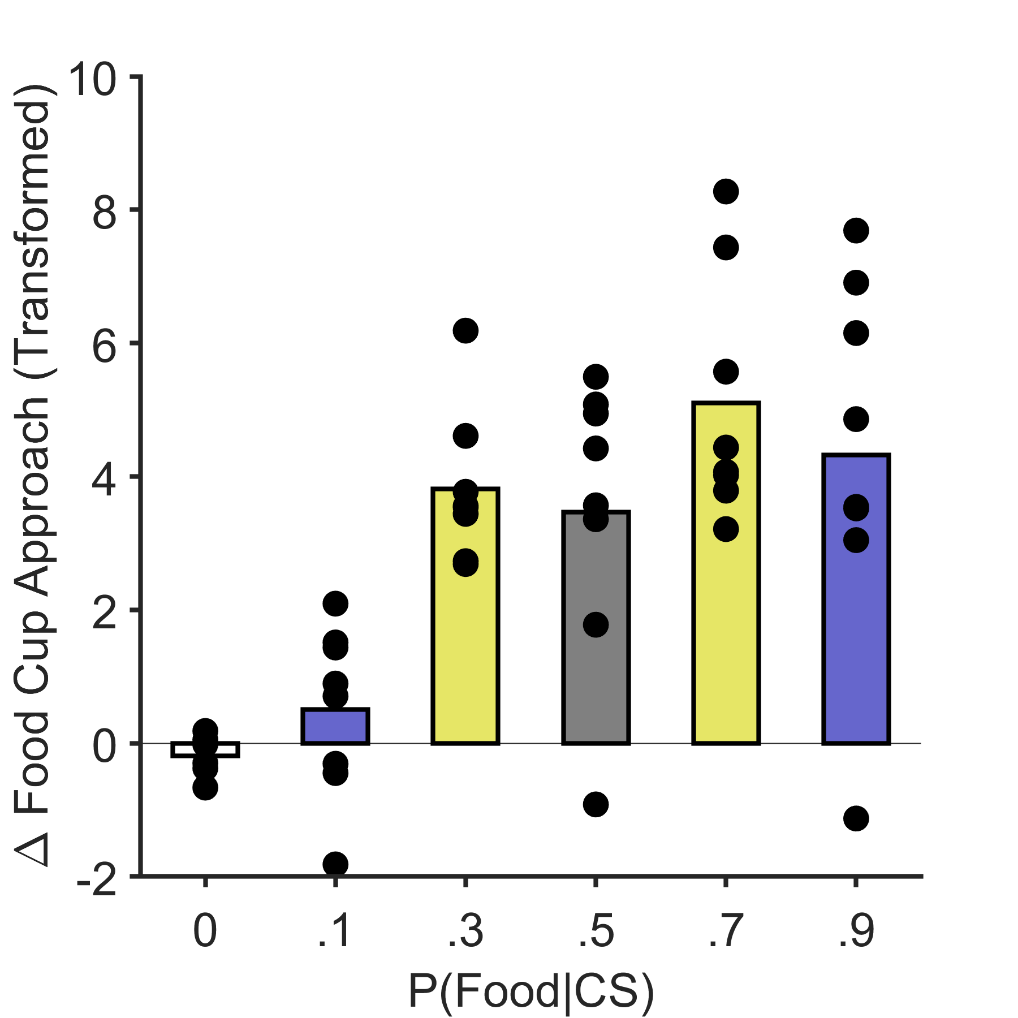

**Supplemental Figure 3**. Pavlovian training in Experiment 1. Cue-induced changes in food-cup approach behavior increased with expected reward probability. Data are averaged over the final 3 days of Pavlovian training. Bars represent the mean rate of food cup approaches (response per minute) during the pre-cue period subtracted from the rate of food cup approaches during the cue period. These data represent the square-root transforms of the data presented in Figure 1A (i.e., the data used in the statistical analysis). Individual rats’ data are indicated by the data points.

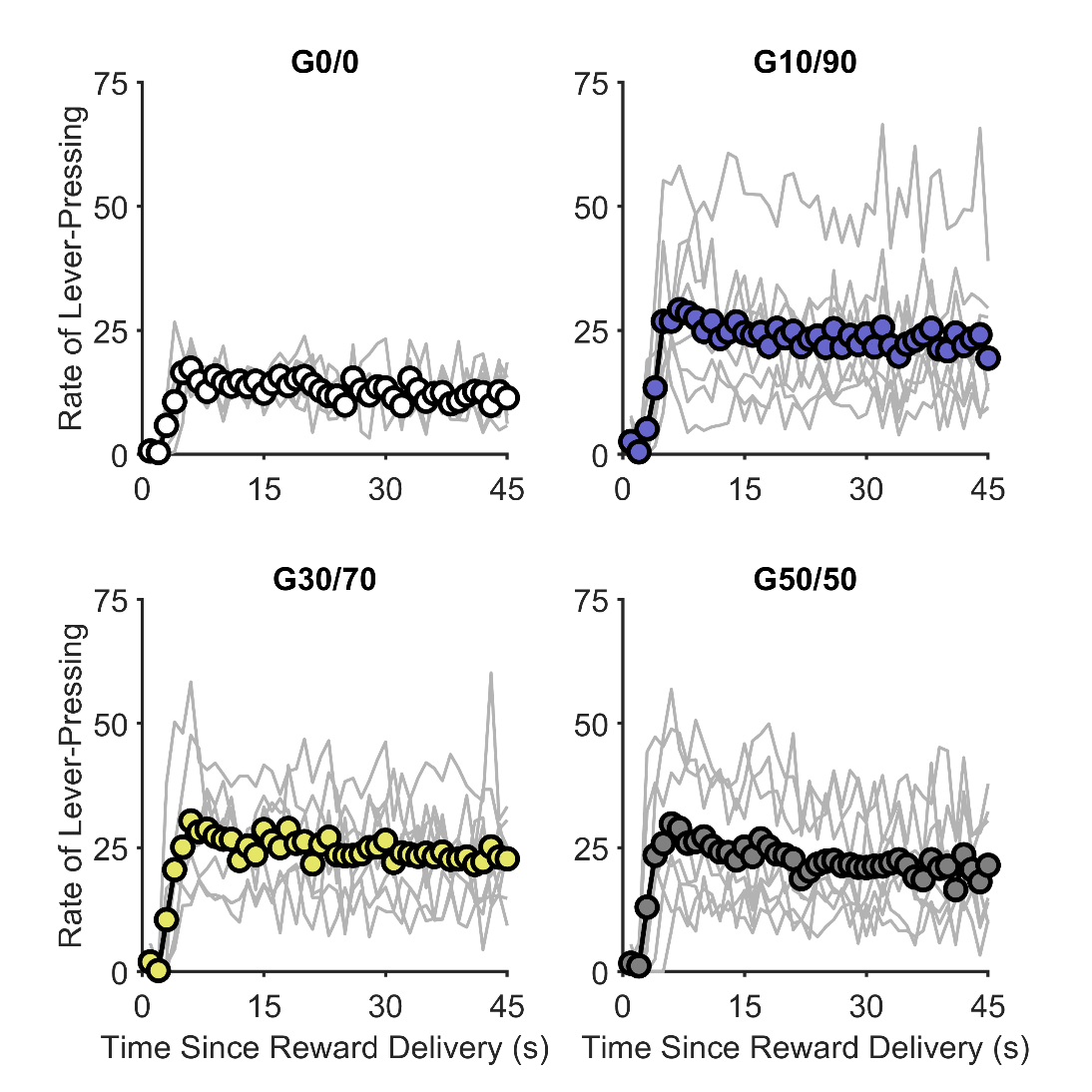

**Supplemental Figure 4**. Response gradients during the final 3 days of instrumental training. Except for Group 0/0 (G0/0), all groups responded at similar rates. Data represent the rate of lever pressing (i.e., responses per minute, controlling for the number of opportunities to respond in each time bin). Group means (circles) are the same as in Figure 1B. Individual rats’ data are indicated by thin lines. G = group.

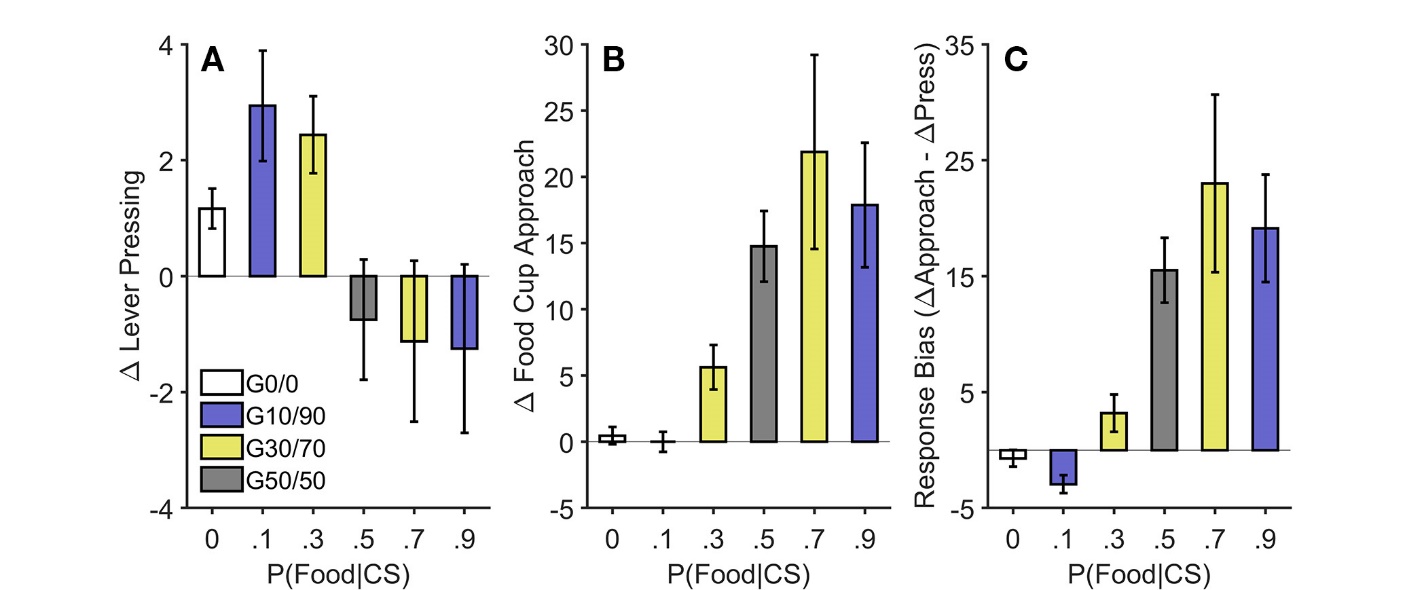

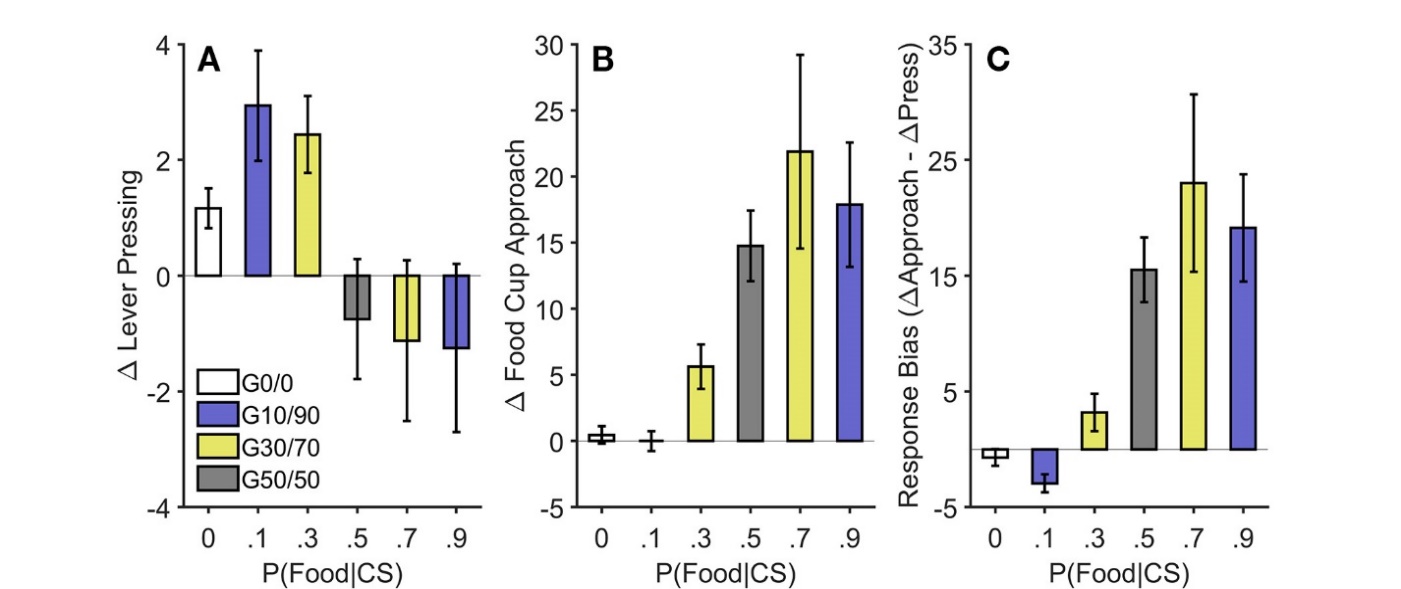

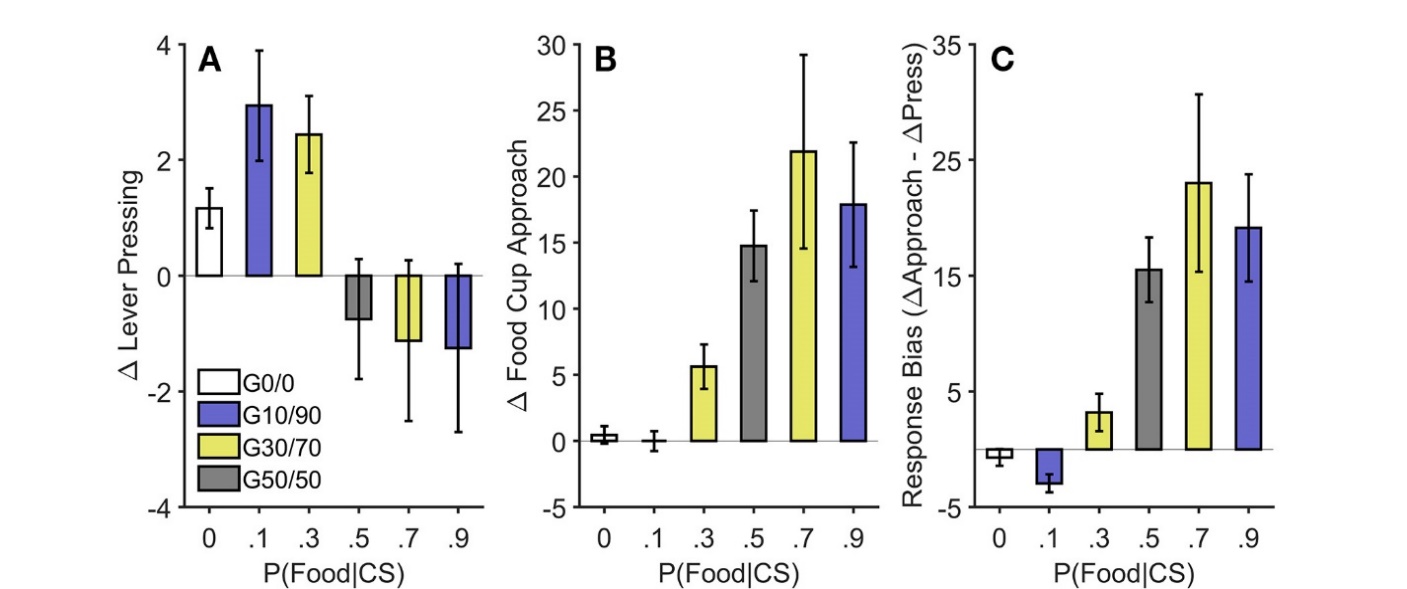

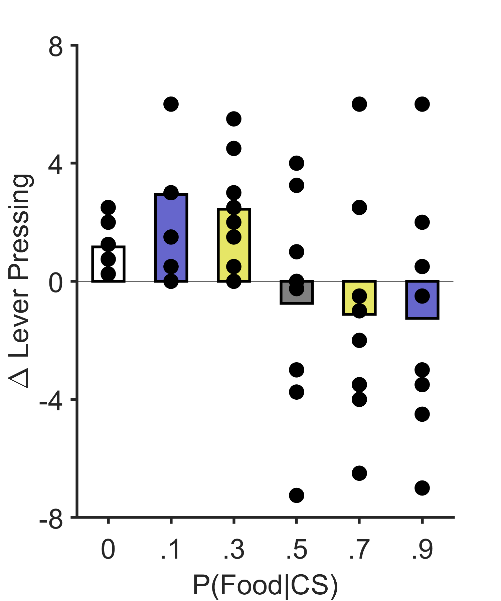

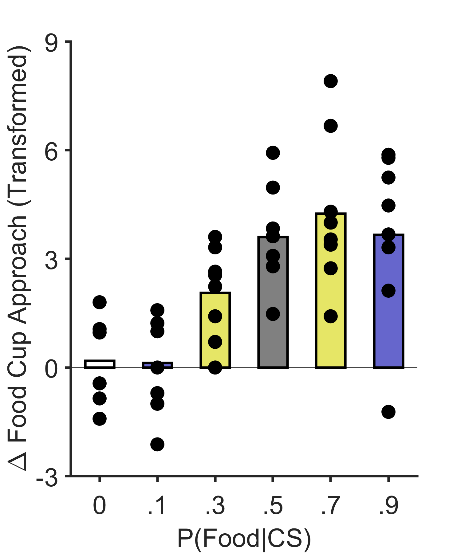

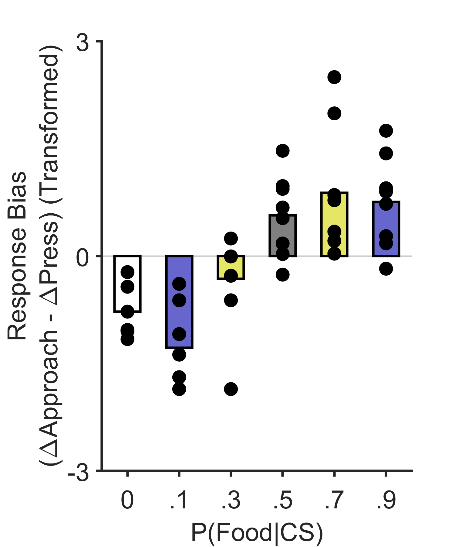

**Supplemental Figure 5**. Pavlovian-to-instrumental transfer in Experiment 1. (A) Cues that signaled a low probability of reward were more effective at eliciting lever pressing than cues signaling a high probability of reward. Data represent the rate of lever pressing (i.e., responses per minute) during the pre-cue period subtracted from the rate of lever pressing during the cue period and 10-s post-cue period. Group means are the same as they are in Figure 2A. Individual rats’ data are indicated by the data points. (B) Concurrent changes in food-cup approach behavior during CS presentations increased with expected reward probability. Data represent the rate of food cup approach (i.e., responses per minute) during the pre-cue period subtracted from the rate of food cup approach during the cue period and 10-s post-cue period. These data represent the square-root transforms of the data presented in Figure 2B (i.e., the data used in the statistical analysis). Individual rats’ data are indicated by the data points. (C) The tendency for cues to bias behavior toward the food cup relative to the lever increased with expected reward probability. Data represent cue-induced changes in lever pressing (A) subtracted from cue-induced changes in food cup approach (B). These data represent the Yeo-Johnson transforms of the data presented in Figure 2C (i.e., the data used in the statistical analysis). Individual rats’ data are indicated by the data points. See Methods for additional details. CS = conditioned stimulus. P = probability.

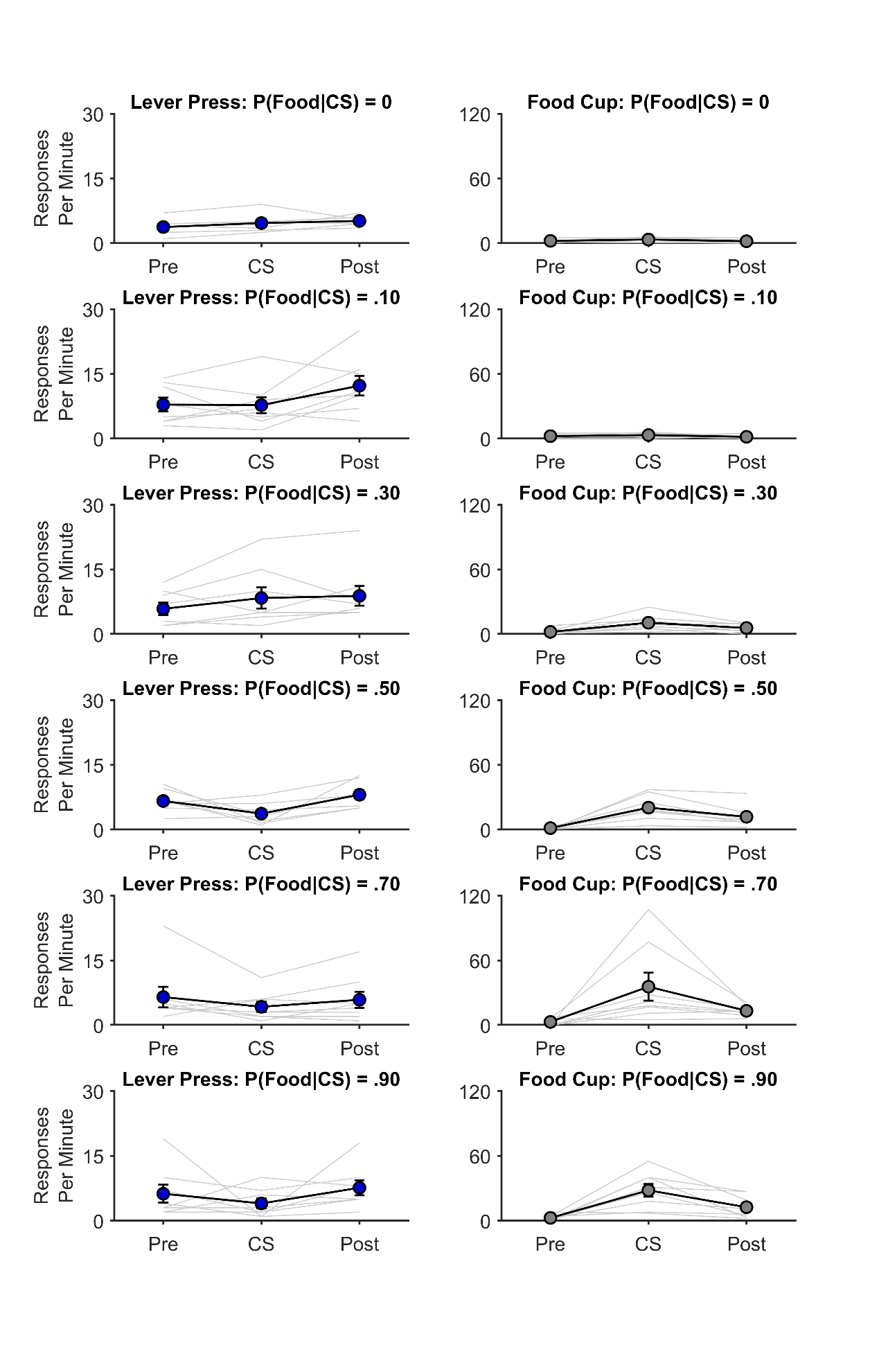

**Supplemental Figure 6**. Response gradients for lever pressing (left panels) and food-cup approach (right panels) in Experiment 1. Data are shown for three 10-s periods, relative to conditioned stimulus (CS) presentation: pre-CS (Pre), during the CS (CS), and post-CS (Post). Thin lines represent individual rats, and data points represent the group means. Error bars reflect ± 1 between-subjects standard error of the mean.

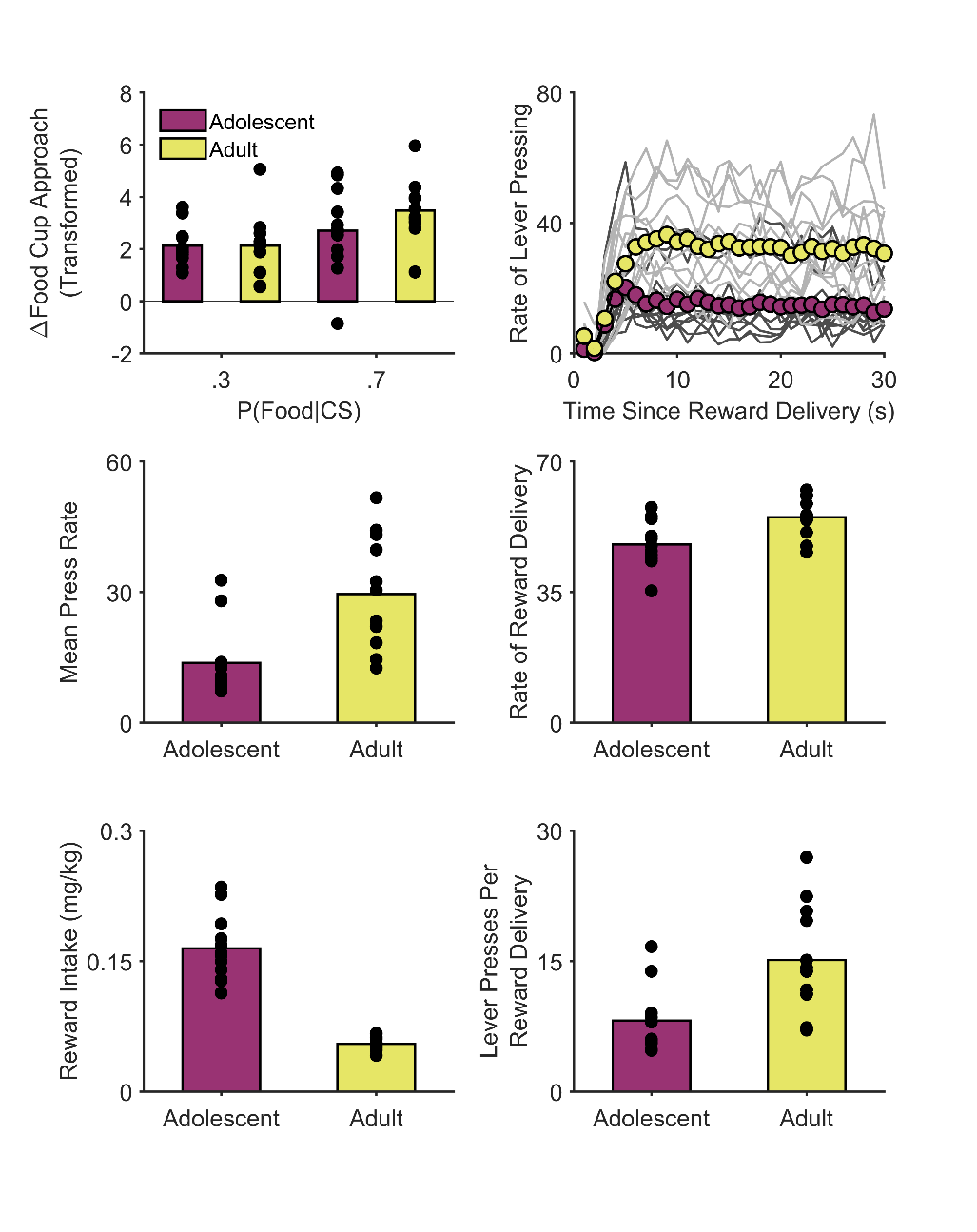

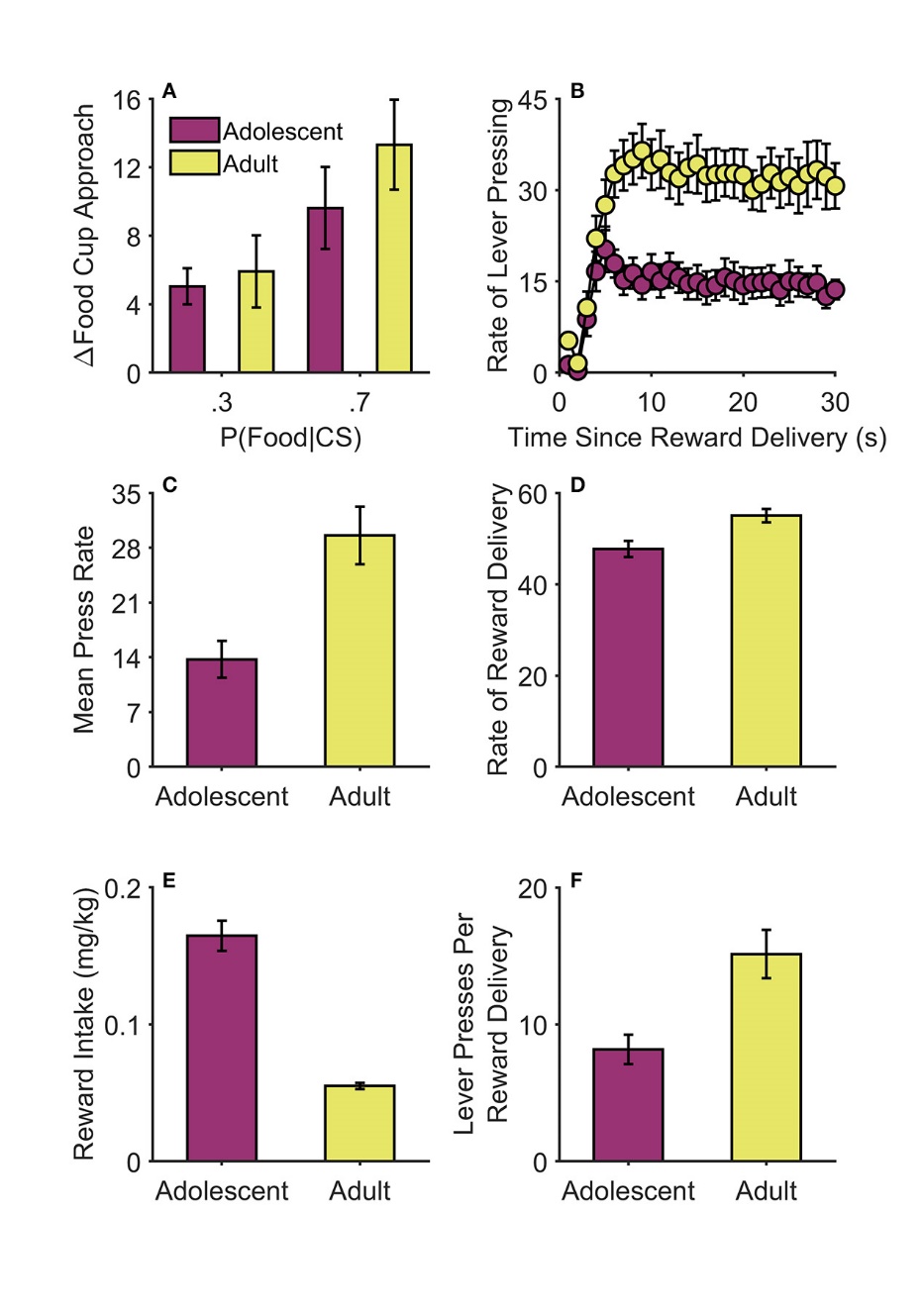

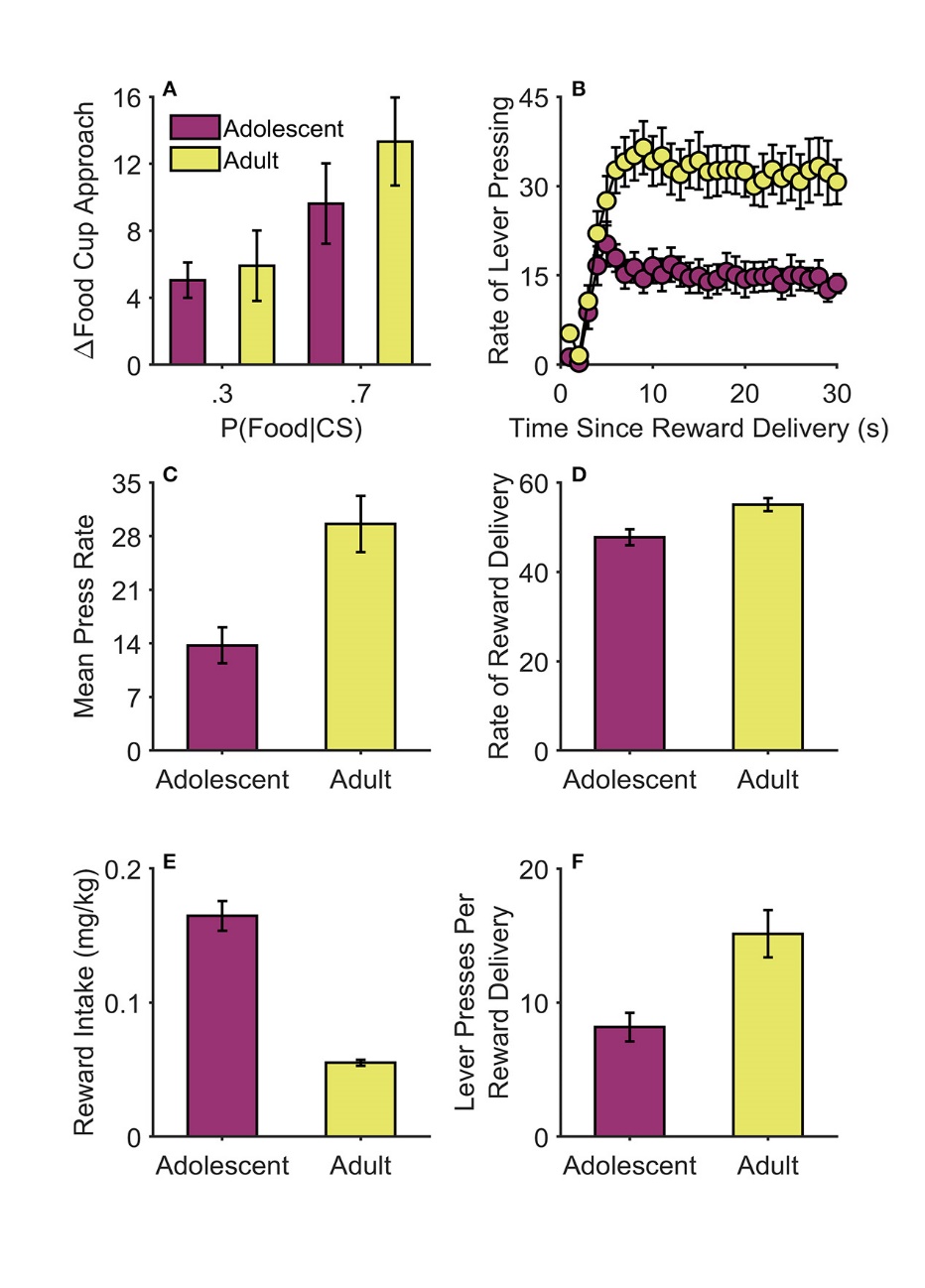

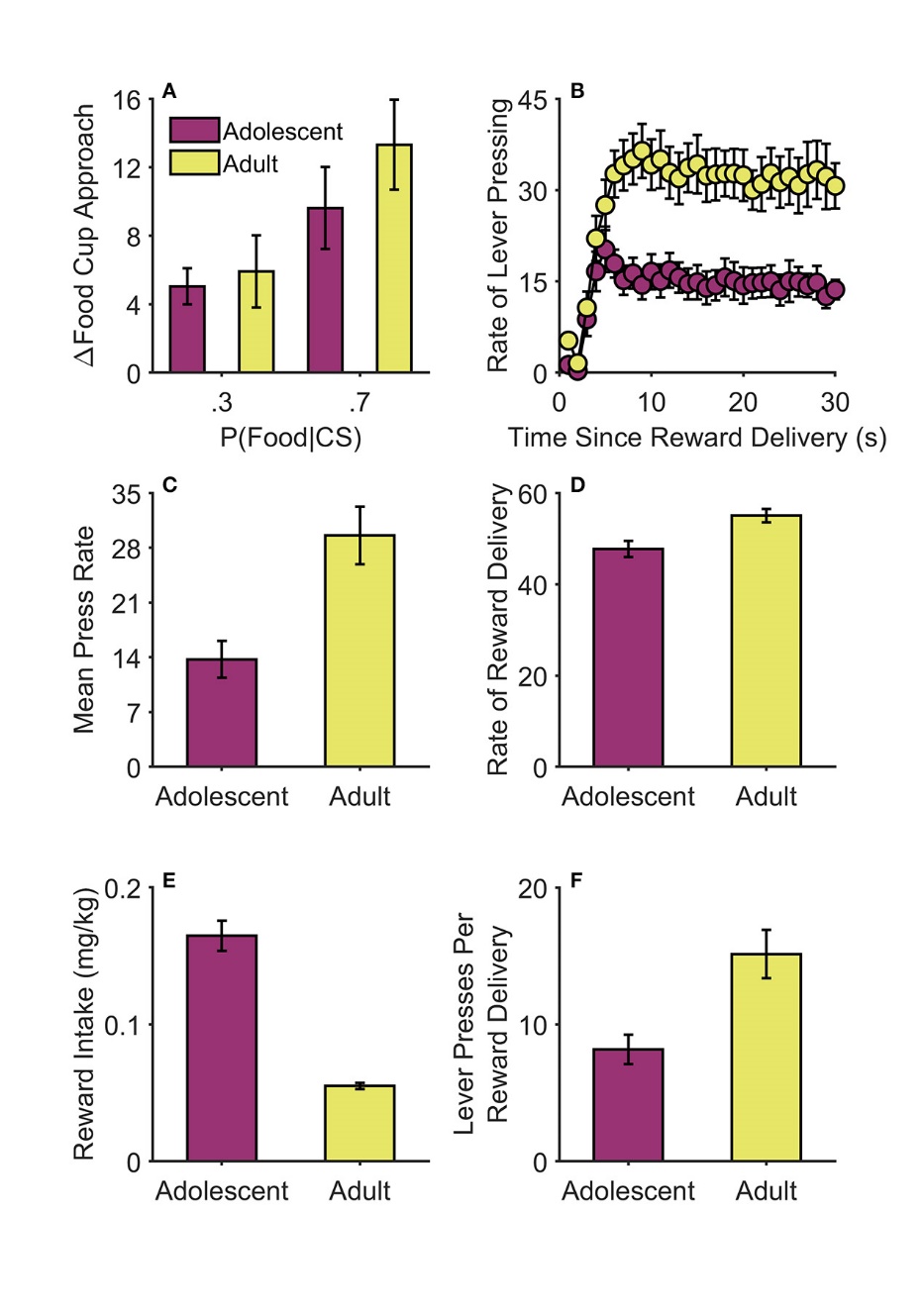

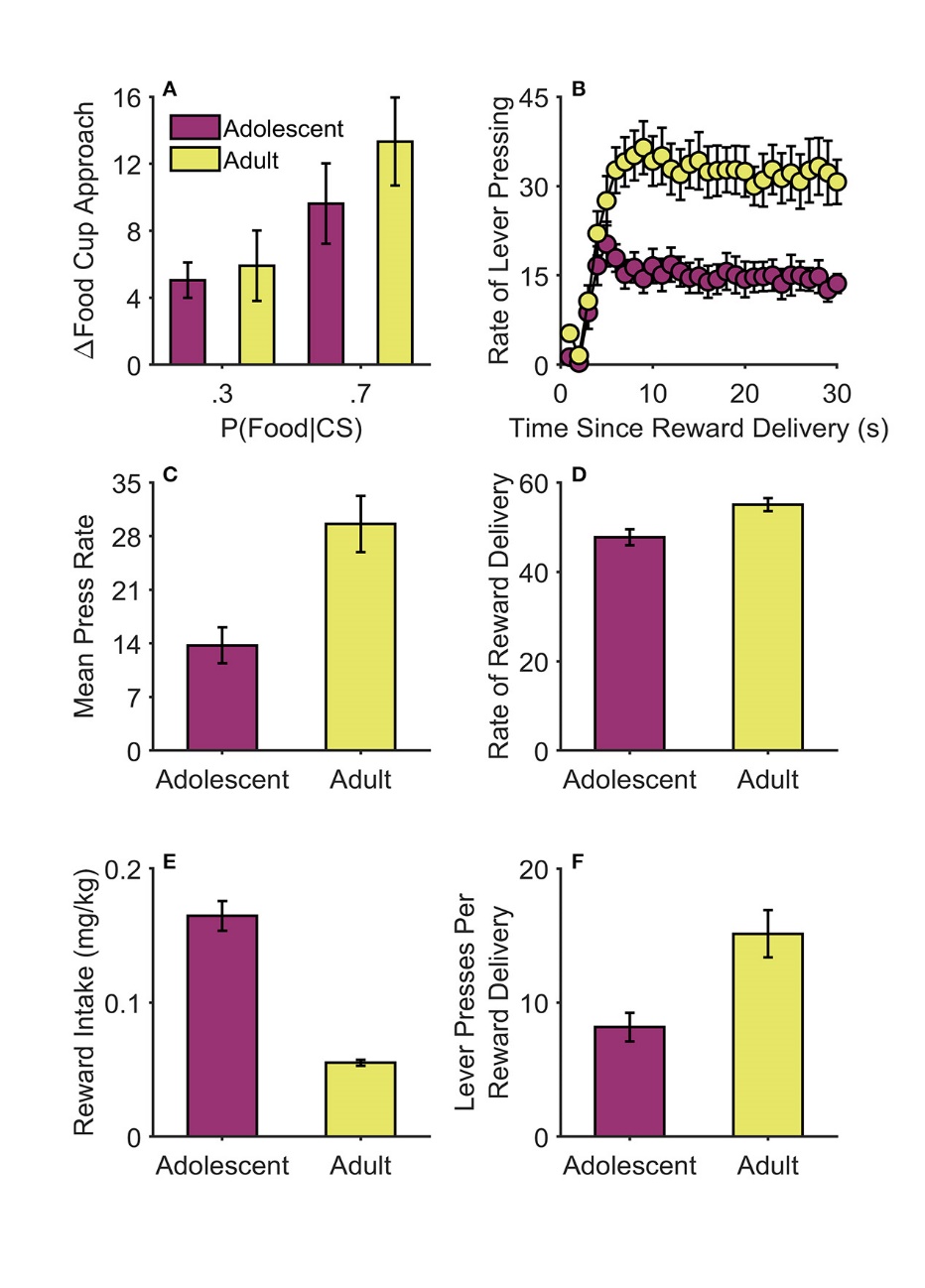

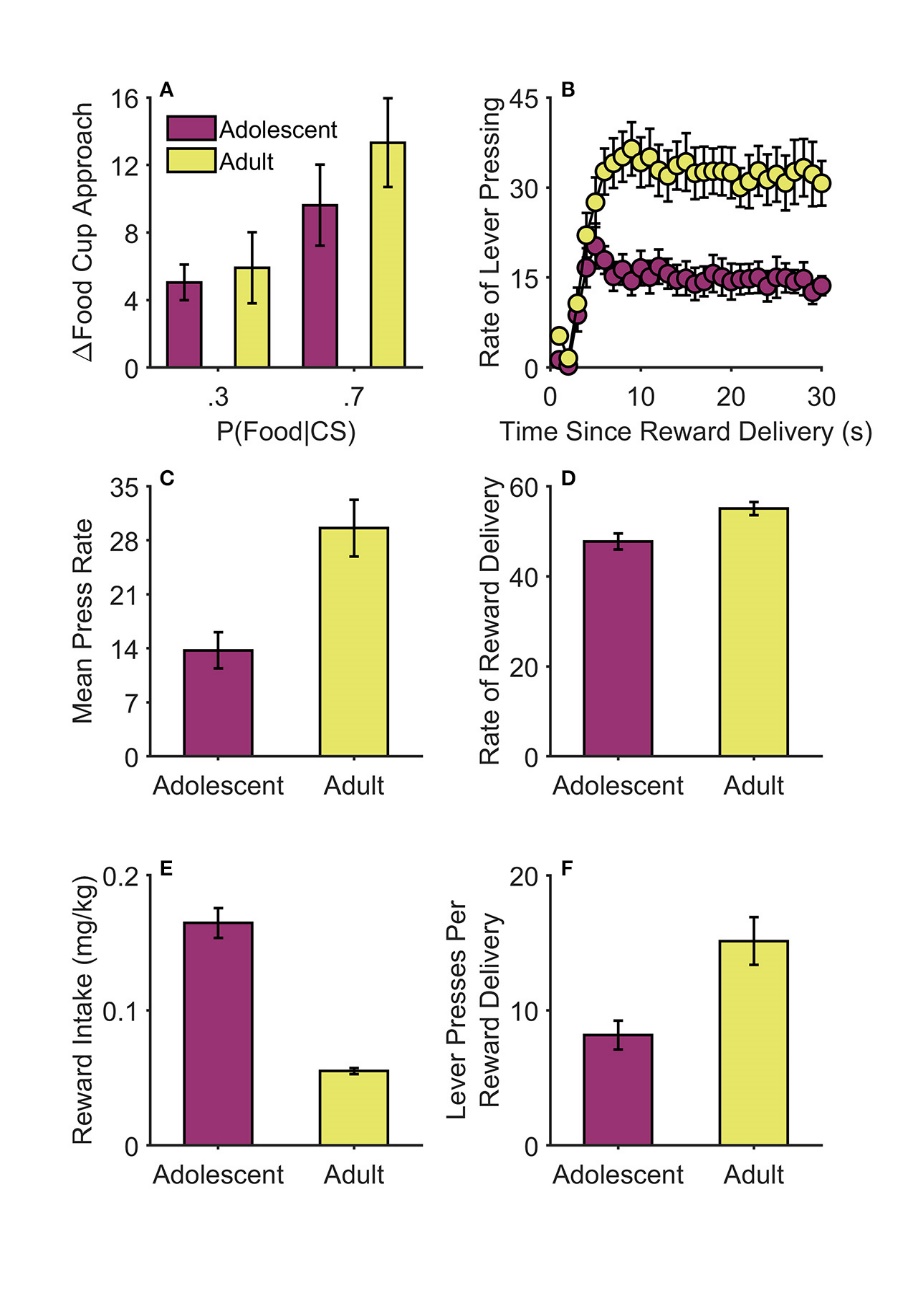

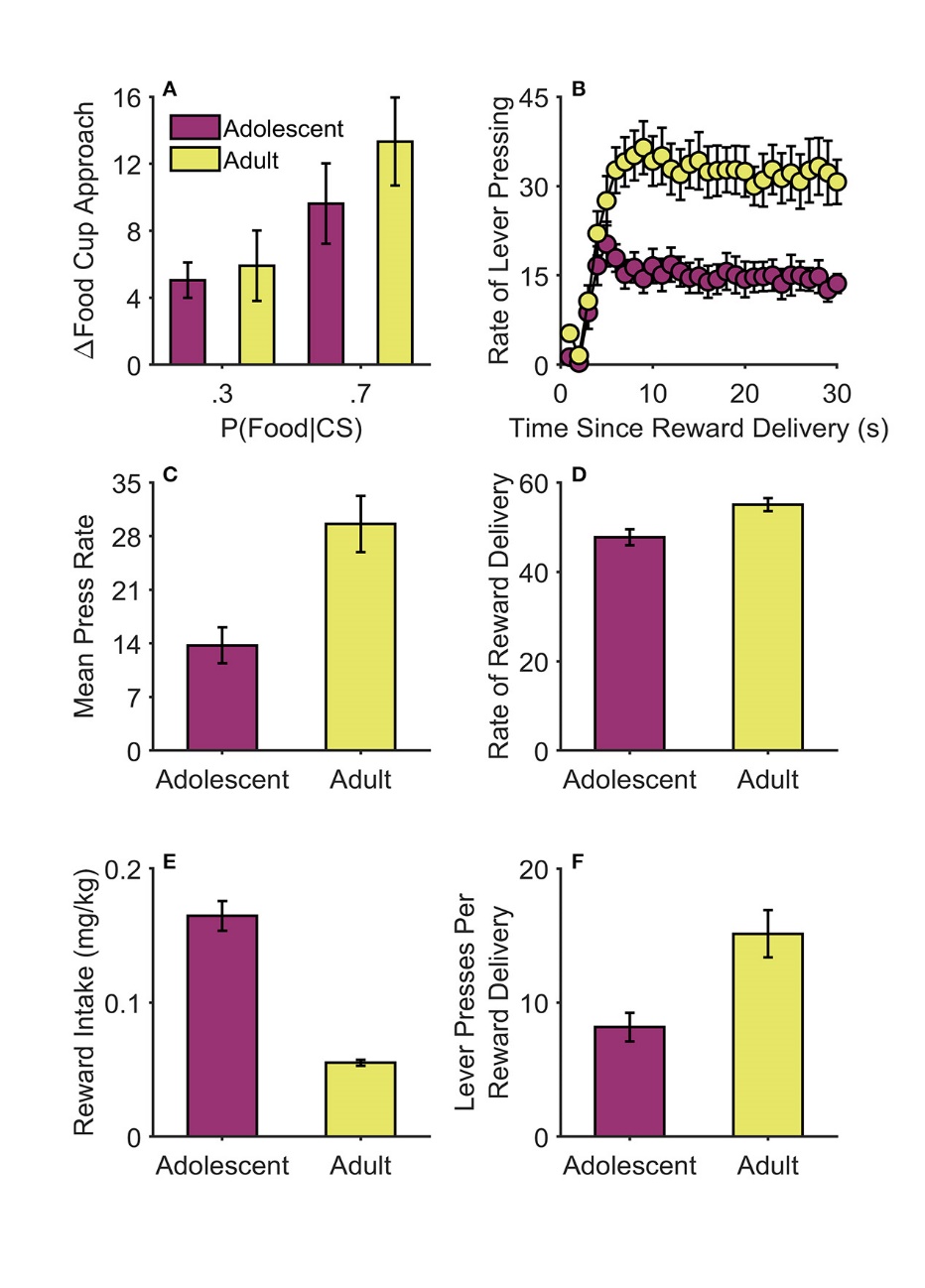

**Supplemental Figure 7**. Pavlovian and instrumental training in Experiment 2. (A) Cue-induced changes in food-cup approach behavior increased with expected reward probability in both adults and adolescents. These data represent the square-root transforms of the data presented in Figure 3A (i.e., the data used in the statistical analysis). (B) Response gradients during the final 3 days of instrumental training. Data represent the rate of lever pressing (i.e., responses per minute, controlling for the number of opportunities to respond in each time bin). Group means (circles) are the same as in Figure 3B. Individual rats’ data are indicated by thin lines (adults are represented by lighter gray lines, adolescents by darker gray lines). (C) Overall, adolescents responded at lower rates than adults. (D) Adults experienced a greater rate of reward delivery than adolescents; data represent mean number of rewards earned per session. (E) In contrast, weight-adjusted reward intake (mg of food pellets per kg of body weight) per session was considerably higher in adolescents. (F) Adolescents were more efficient in their responding than adults, exhibiting fewer lever presses per reward delivery. In Panels A, C, D, E, and F, individual rats’ data are indicated by the data points. See Methods for additional details. CS = conditioned stimulus. P = probability.

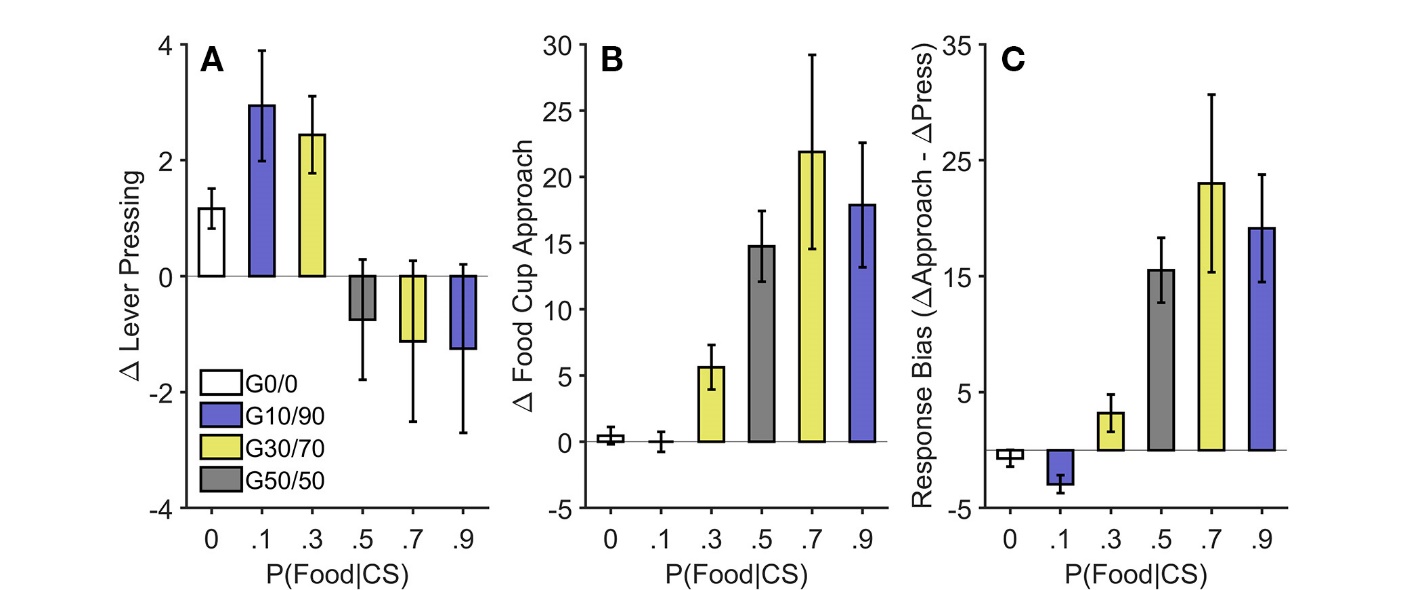

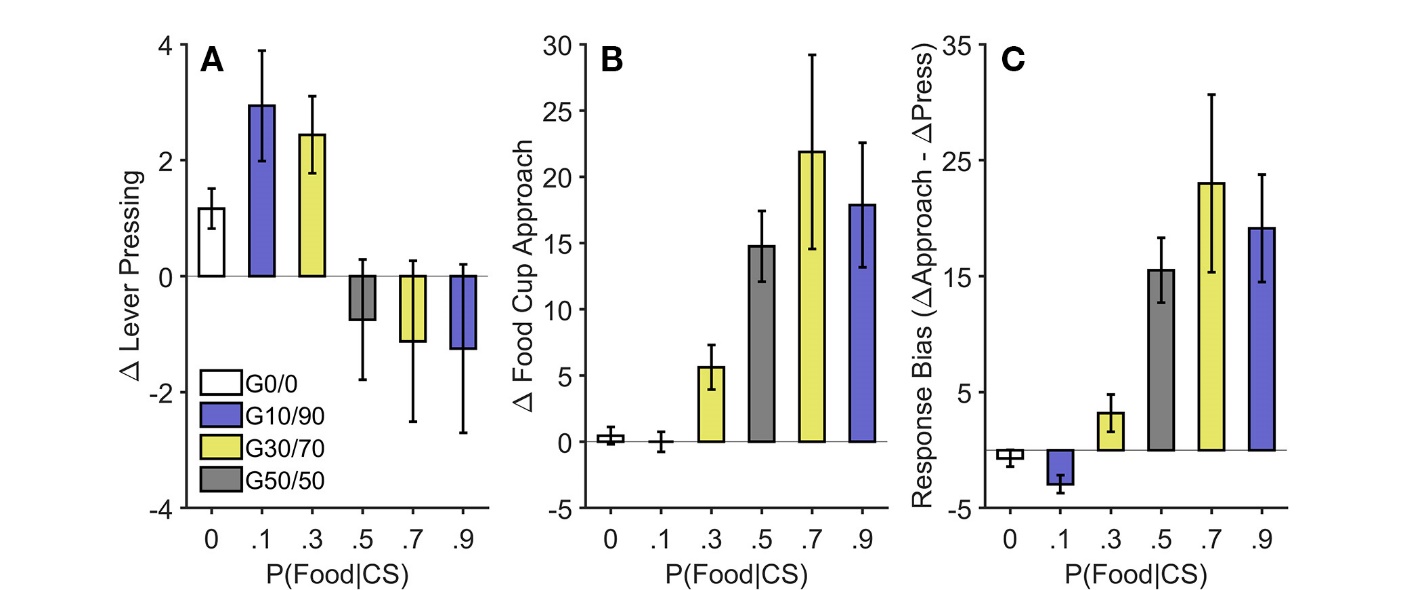

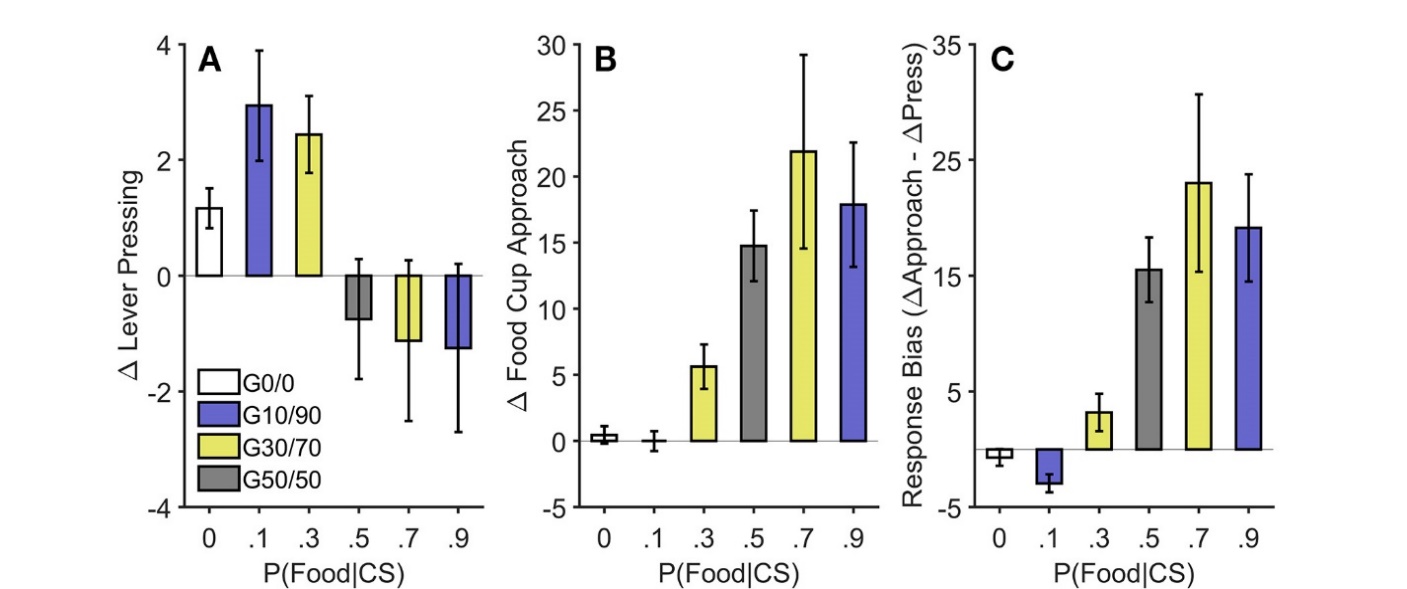

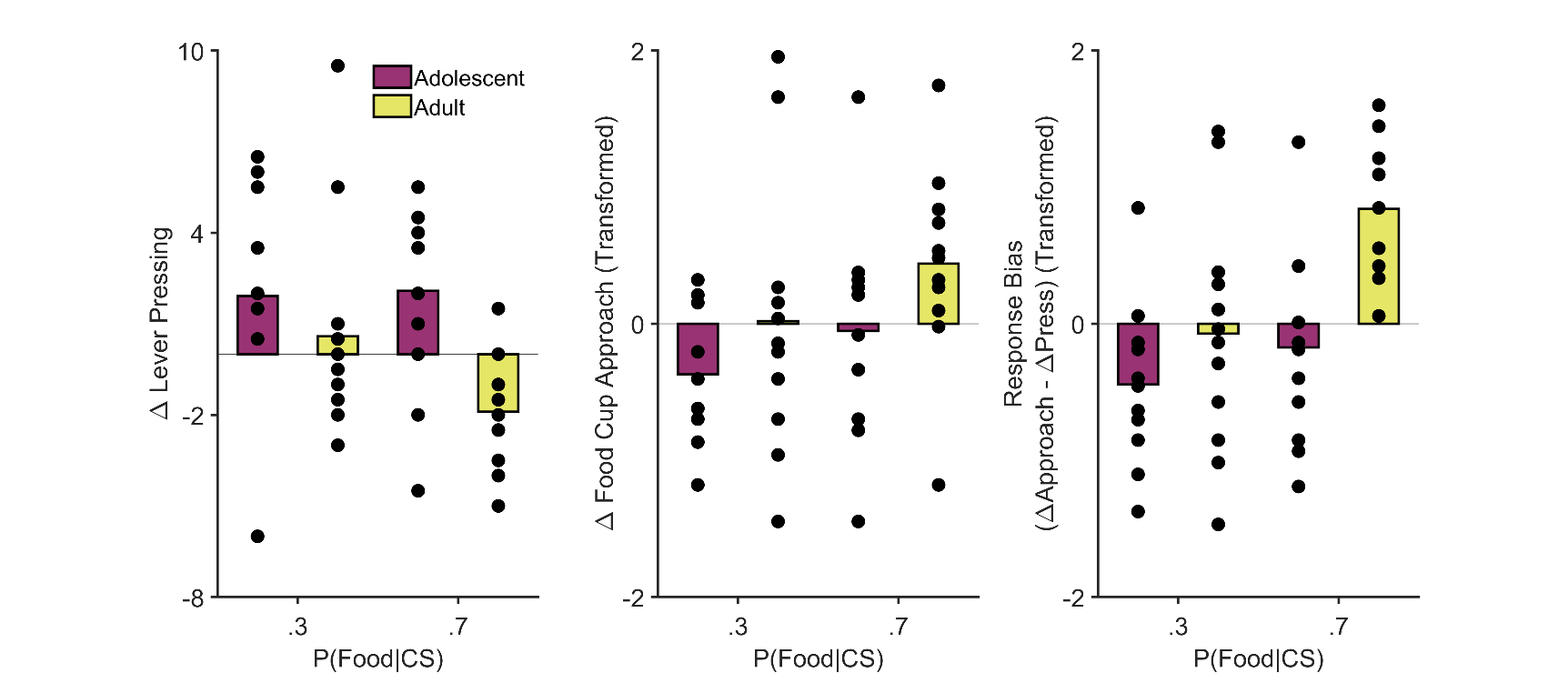

**Supplemental Figure 8**. Pavlovian-to-instrumental transfer in Experiment 2. (A) For adult rats, cue-elicited lever pressing was greater during trials with the 30% CS than the 70% CS. In contrast, adolescent rats showed a similar increase in lever pressing to both cues. Group means are the same as they are in Figure 4A. Individual rats’ data are indicated by the data points. (B) Both groups showed similar patterns of cue-elicited food-cup approach behavior, though adolescent rats showed a marginally lower rate of conditioned food-cup approach. These data represent the Yeo-Johnson transforms of the data presented in Figure 4B (i.e., the data used in the statistical analysis). Individual rats’ data are indicated by the data points. (C) The tendency for cues to bias behavior toward the food cup relative to the lever was greater for adults than for adolescents, particularly during the 70% CS. These data represent the Yeo-Johnson transforms of the data presented in Figure 4C (i.e., the data used in the statistical analysis). Individual rats’ data are indicated by the data points. See Methods for additional details. Error bars reflect ± 1 between-subjects standard error of the mean. CS = conditioned stimulus. P = probability.

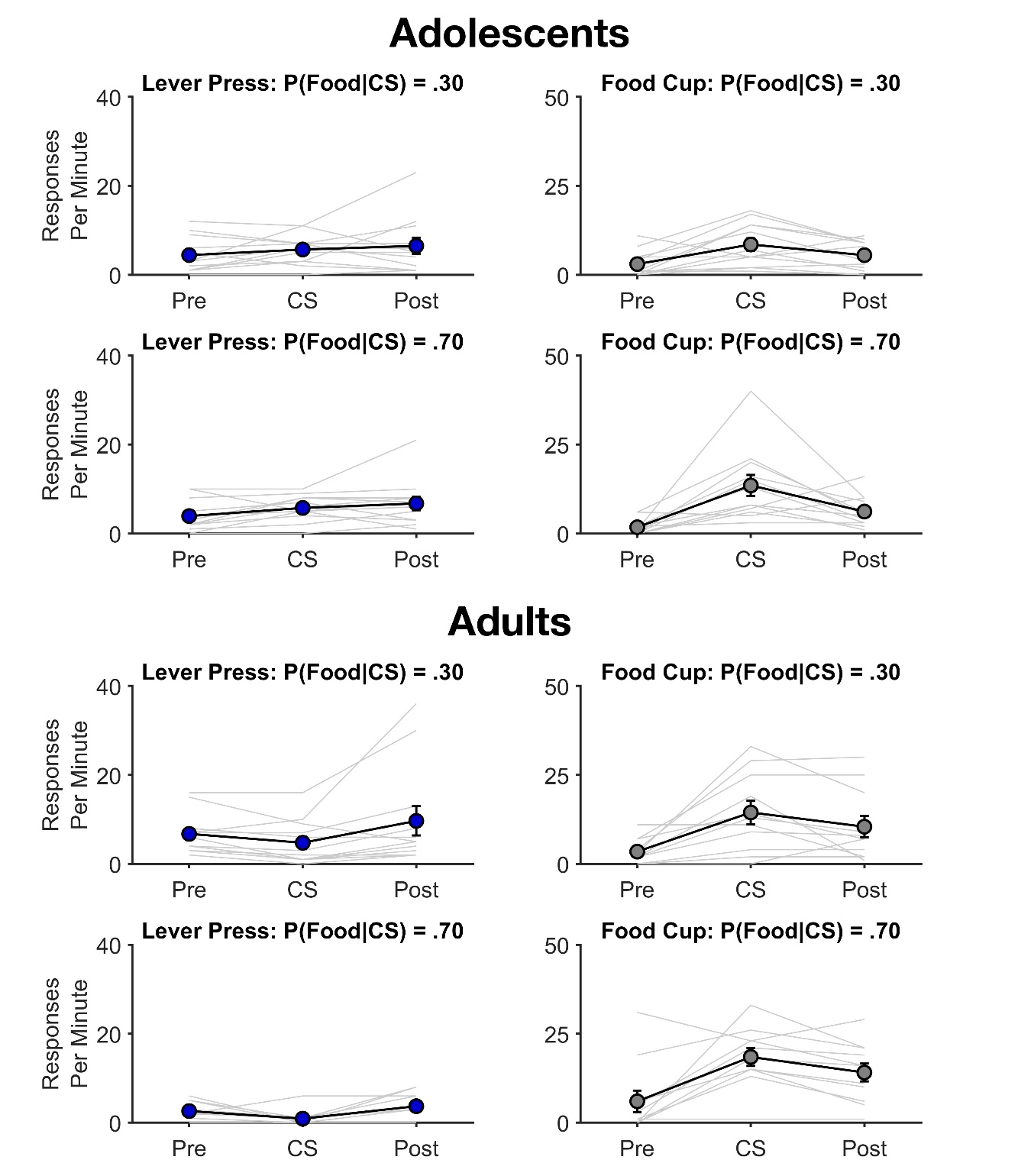

**Supplemental Figure 9**. Response gradients for lever pressing (left panels) and food-cup approach (right panels) in Experiment 2. Data are shown for three 10-s periods, relative to conditioned stimulus (CS) presentation: pre-CS (Pre), during the CS (CS), and post-CS (Post). Thin lines represent individual rats, and thick lines and data points represent the group means. Error bars reflect ± 1 between-subjects standard error of the mean.
